# Integrative Epigenomic and Transcriptomic Analysis Reveals Robust Metabolic Switching in the Brain During Intermittent Fasting

**DOI:** 10.1101/2020.03.22.002725

**Authors:** Gavin Yong-Quan Ng, Dominic Paul Lee Kok Sheng, Sung Wook Kang, David Yang-Wei Fann, Joonki Kim, Asfa Alli-Shaik, Yoon Suk Cho, Jihoon Han, Jae Hoon Sul, Vardan Karamyan, Eitan Okun, Thameem Dheen, M. Prakash Hande, Raghu Vemuganti, Karthik Mallilankaraman, Brian K. Kennedy, Grant Drummond, Christopher G. Sobey, Jayantha Gunaratne, Mark P. Mattson, Roger Sik-Yin Foo, Dong-Gyu Jo, Thiruma V. Arumugam

## Abstract

Intermittent fasting (IF) is a lifestyle intervention comprising a dietary regimen in which energy intake is restricted via alternating periods of fasting and *ad libitum* food consumption, without compromising nutritional composition. While epigenetic modifications can mediate effects of environmental factors on gene expression, no information is yet available on potential effects of IF on the epigenome. In this study, we found that IF causes modulation of histone H3 lysine 9 trimethylation (H3K9me_3_) epigenetic mark in the cerebellum of male C57/BL6 mice, which in turn orchestrates a plethora of transcriptomic changes involved in the robust metabolic switching processes commonly observed during IF. Interestingly, both epigenomic and transcriptomic modulation continued to be observed after refeeding, suggesting that memory of the IF-induced epigenetic change is maintained at the locus. Notably though, we found that termination of IF results in a loss of H3K9me_3_ regulation of the transcriptome. Collectively, our study characterizes a novel mechanism of IF in the epigenetic-transcriptomic axis, which controls myriad metabolic process changes. In addition to providing a valuable and innovative resource, our systemic analyses reveal molecular framework for understanding how IF impacts the metaboloepigenetics axis of the brain.

**Highlights:**

○ Intermittent fasting (IF) and refeeding modifies epigenome in the cerebellum
○ Integrative epigenomic and transcriptomic analyses revealed metabolic switching
○ IF affects the metaboloepigenetics axis in regulating metabolic processes
○ Integrative analyses revealed a loss of epigenetic reprogramme following refeeding

## 1. Introduction

Intermittent fasting (IF) is a dietary regimen that restricts energy intake by alternate periods of fasting and *ad libitum* food consumption, without compromising nutritional composition. Many animal studies have established that IF ameliorates the development of age-related cardiovascular, neurodegenerative and metabolic diseases and promotes longevity. Despite extensive experimental evidence supporting such protection by IF, our understanding of the underlying molecular mechanisms is still poor. Epigenetic modifications have been shown to be pivotal in mediating the influence of environmental factors on genomic status. Indeed, many age-related diseases are polygenic and confounded by environmental influences (Boyle et al., 2015), suggesting that interactions between the environment and genetic framework may underlie the complex pathophysiology of chronic diseases.

Calorie restriction (CR) is one form of environmental manipulation that can impact the epigenome through various epigenetic modifications and consequently affect a plethora of biological pathways that modulate age-related epigenetic events. For instance, CR can influence the epigenetic regulation of immuno-metabolic adaptation and attenuate age-dependent epigenetic drift, thus reducing the pathogenesis of age-related diseases (Hernández-Saavedra et al., 2019; Maegawa et al., 2017; Molina-Serrano and Kirmizis, 2017). IF is a lifestyle intervention representing a potential environmental factor capable of influencing an individual’s epigenome. However, understanding of IF-induced epigenetic changes is currently lacking and the potential epigenetic effects of refeeding after IF are unknown. Thus, it is of great interest whether IF may serve as an environmental trigger that can influence epigenomic profile.

H3K9me_3_ is often associated with constitutive heterochromatin or inactive euchromatin, and has been extensively reported to be modulated during CR (Molina-Serrano et al., 2019; Sen et al., 2016; Vaquero and Reinberg, 2009; Xie et al., 2016). In studies using the epigenetic clock (Horvath et al., 2015), the cerebellum, a major brain region harbouring a large population of neuronal cells (Bahney et al., 2017; WallÃ̧e et al., 2014), was recently reported to age more slowly than other parts of the human body, and to contain a circadian oscillator that responds robustly to energy restriction (Delezie et al., 2016; Mendoza et al., 2010). It was hypothesized that as the large population of cells within the cerebellum would require high energy demand, they should be sensitive to energy restriction and induce robust metabolic changes in response. Thus, we decided to investigate the effects of IF and refeeding on the modulation of H3K9me_3_ in the cerebellum. We studied the epigenomic and transcriptomic profiles resulting from two common IF regimens, time-restricted fasting for 16 hours (IF16) or 24 h on alternate days (i.e. ‘every other day’; EOD), in mice for three months. Also, we assessed changes in epigenomic and transcriptomic patterns after refeeding in mice refed for two months (IF16.R or EOD.R) following the IF regimens. We found that both types of IF regimen were capable of distinctly affecting the modulation of H3K9me_3_, which in turn differentially regulated different aspects of the transcriptome, especially in the metabolic axis. Moreover, we found that following refeeding, mice that had been subjected to either IF regimen demonstrated differential epigenomic and transcriptomic profiles. However, following abolition of IF, there was a loss of epigenetic memory in the transcriptome. Our data thus provide novel insights into the epigenetic-transcriptomic axis of IF and refeeding mechanisms in the cerebellum.

## 2. Results

### 2.1 IF mice show decreased body weight without a change in fuel preferences

The summarized study design includes the timing of interventions and blood and tissue collection **(Figure 1)**. Male C57BL/6N mice were fed a normal chow diet (on a caloric basis: 58%, 24%, and 18% of carbohydrate, protein, and fat, respectively). Mice were randomly assigned to one of the three study groups, AL (*ad libitum*), daily IF16, or EOD schedules, beginning at 3 months of age. First, we monitored the physiological effect of IF and the refeeding regimens on C57/BL6 mice. Both IF16 and EOD groups had a lower body weight than AL mice after three-months of IF (**Figure S1a**), but were not different from each other. After the two-months refeeding regimen, i.e. IF16.R, EOD.R, and AL.R mice, did not differ in body weight (**Figure S1a**). Aging in mice was associated with an increase in their average body weight (**Figure S1a**). Overall, the energy intake of mice subjected to IF or refeeding was not different to that of control mice (**Figure S1b**). Next, we investigated the composition of energy intake to examine whether mice compensated for energy restriction through a preferential shift in the specific macromolecules from which energy was obtained. Notably, the energy intake from each compositional source (carbohydrate, fat, protein) was not different across the three study groups (AL, IF, EOD) (**Figure S1c**). However, mice in the corresponding refeeding group (AL.R, IF.R, EOD.R) consistently showed a higher energy intake for each compositional source (**Figure S1c**). Our findings, therefore exclude the possibility that differences in body weight during IF were due to an altered composition of energy intake.

**Figure 1:**
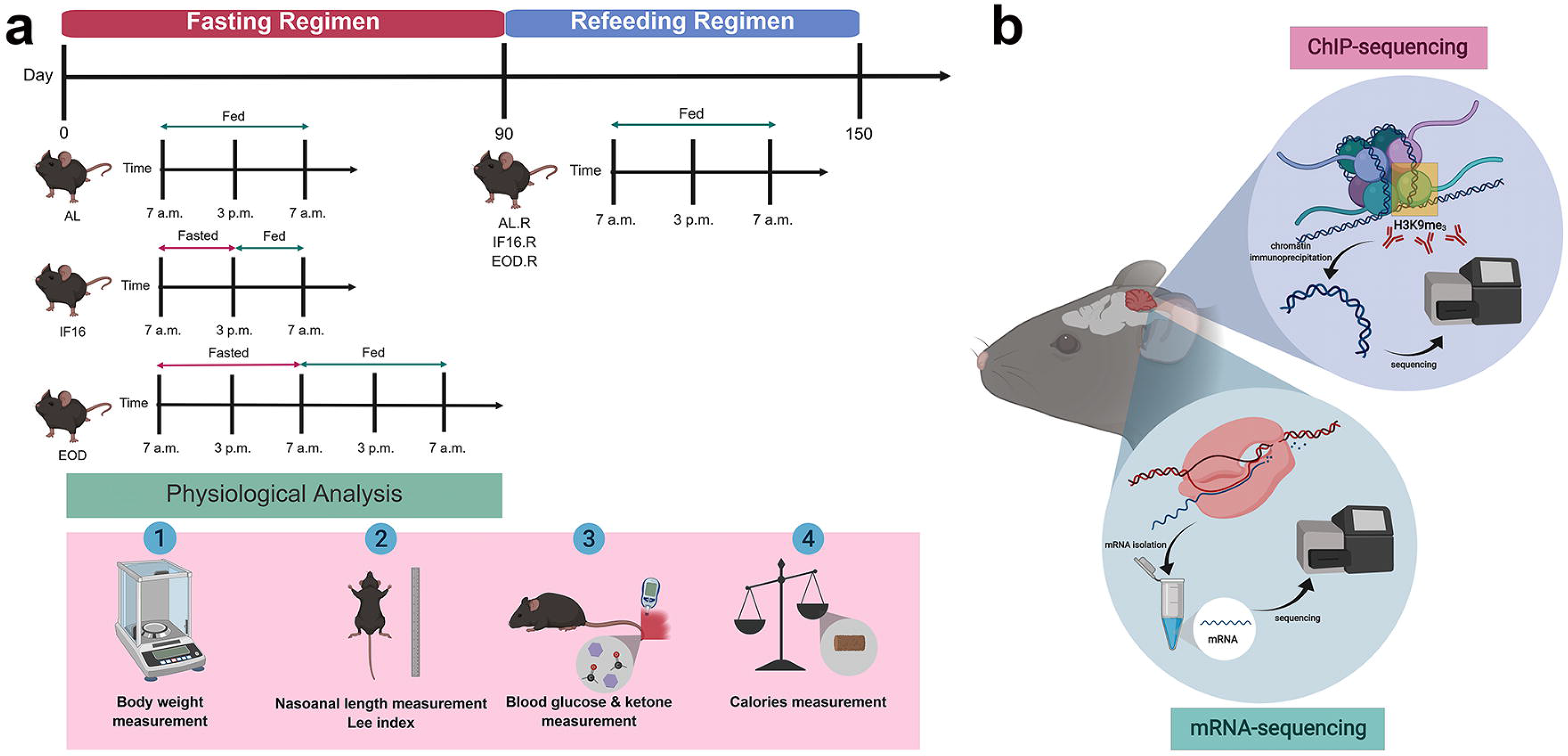
Experimental design for intermittent fasting and refeeding regimen. (a) Mice were raised to 3 months-old with *ad libitum* access to food before being randomly assigned and subjected to AL feeding, IF16 or EOD for a further 3 months. All mice had AL access to water, with AL mice having free access to food also. During the refeeding regimen, the three groups of mice were allowed AL access to both food and water for a further 2 months. A series of physiological tests was performed on the mice as depicted. (b) Cerebellar tissue was harvested, frozen and subjected to chromatin immunoprecipitation at H3K9me_3_ locus and eukaryotic mRNA extraction prior to sequencing via Illumina platforms. AL, ad libitum; IF16, intermittent fasting 16 hours; EOD, intermittent fasting 24 hours on alternate day or ‘every other day’; AL.R, AL and refeed; IF16.R, IF16 and refeed; EOD.R, EOD and refeed; H3K9me_3_, histone 3 lysine 9 trimethylation.

Blood analyses showed that both IF16 and EOD mice had lower glucose and higher ketones than AL mice (**Figure S2a and b**). Interestingly, prolonged energy restriction resulted in a greater change in both glucose and ketone homeostasis, suggesting that the differential impact of energy restriction on fuel utilization is time-dependent. However, following refeeding, no differences were observed in blood glucose levels across study groups but a significant change in blood ketone levels was observed in EOD.R mice. (**Figure S2a and b**). Concordantly, body composition analysis indicated a reduction in the Lee index or the body fat mass, in both IF16 and EOD mice, as compared to AL mice (**Figure S2c**). No differences were observed in body fat mass following refeeding (**Figure S2c**). Thus, our data show that IF and refeeding have a differential impact on physiological homeostasis in mice.

### 2.2 IF induces epigenetic modifications in the cerebellum that are maintained following refeeding

To establish whether IF may influence the epigenetic landscape, we carried out ChIP (chromatin immunoprecipitation) of the H3K9me_3_ locus in the cerebellum and sequenced the resulting fragments by next-generation sequencing (NGS). ChIP-seq data from the three study groups, AL, IF16, and EOD, were compared on a three-dimensional principal component analysis (3D-PCA) score plot (**Figure 2a**). The 3D-PCA plot showed differential occupancy sites for IF16 and EOD as compared to AL, suggesting that the gene expression patterns modulated by H3K9me_3_ may be distinct between the two IF regimens. The profiles of ChIP-seq data for both, IF16 and EOD, at the H3K9me_3_ mark around transcriptional start sites (TSSs) of annotated genes were different than AL (**Figure 2b**). Gene expression at the H3K9me_3_ locus was enriched in IF16 relative to AL, whereas it appeared to be repressed in EOD mice. Comparision of peaks (significantly enriched genomic intervals in the ChIP-seq dataset) profiles identified 939 differential peaks in IF16 and 241 differential peaks in EOD, as compared to AL, respectively (**Figure 2c**). Of these, 123 differential peaks were common between IF16 and EOD (**Figure 2c**). Collectively, our observations demonstrate that IF modulates the epigentetic landscape in the cerebellum by inducing differential H3K9me_3_ mark on several genes with some being conserved across both the IF regimens and others distinct for either IF16 or EOD groups.

**Figure 2:**
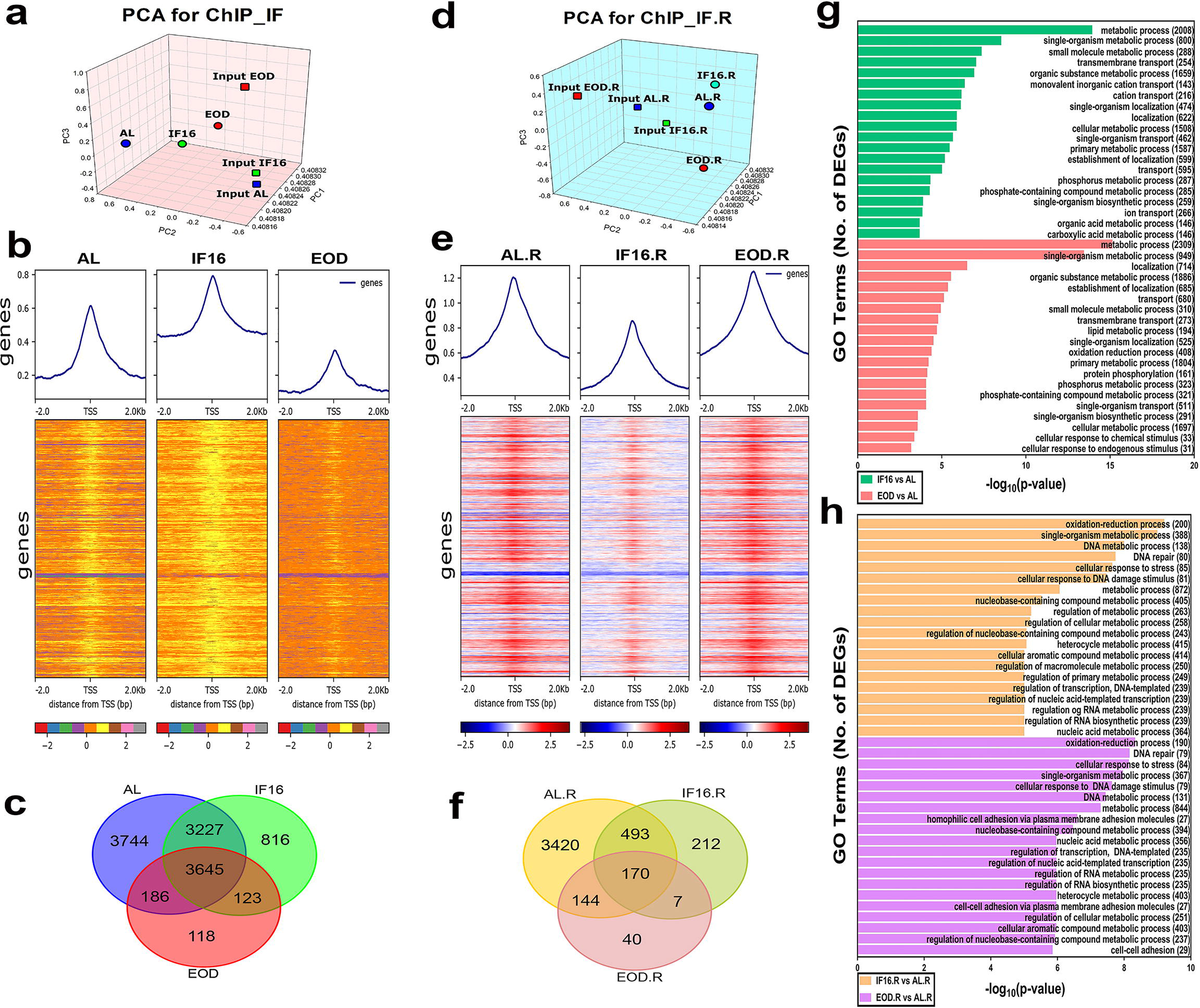
Epigenomic analysis of IF and refeeding at the H3K9me_3_ locus in the cerebellum. (a) Three-dimensional principal component analysis (3D-PCA) plot of H3K9me_3_ ChIP dataset from expression profiles of IF groups. The three most significant principal components (PC1, PC2, and PC3) are displayed on the x-, y-, and z-axes, respectively. PCA discriminated AL, IF16, and EOD into three unique cluster regions relative to their input control. (b) Summary and heatmap plots displaying H3K9me_3_ ChIP-seq signal mapping to a 2 kb window around the TSS of genes revealed distinct expression patterns in AL, IF16, and EOD. ChIP-seq signals are sorted according to mean score and scale bar illustrate log_2_ ratio of ChIP signal vs control signal. (c) Venn diagram illustrates the number of differentially expressed peaks that are common and distinct in each IF regimen when compared to AL. (d) 3D-PCA plot of H3K9me_3_ ChIP dataset obtained from expression profiles of refeeding groups. The three most significant principal components (PC1, PC2, & PC3) are displayed on the x-, y-, and z-axes, respectively. PCA discriminated AL.R, IF16.R, and EOD.R into three unique cluster regions relative to their input control. (e) Summary and heat map plots displaying H3K9me_3_ ChIP-seq signal mapping to a 2 kb window around the TSS of genes revealed distinct expression patterns in AL.R, IF16.R and EOD.R. ChIP-seq signal over the body of genes are sorted according to mean score and scale bar illustrate log_2_ ratio of ChIP signal vs control signal. (f) Venn diagram illustrates the number of differentially expressed peaks that are common and distinct in each refeeding regimen when compared to AL.R. (g-h) Top 20 differentially expressed peaks-associated gene ontologies of IF and refeeding compared to control plotted against statistical significance (represented as (−log_10_ p-value). Number of differentially expressed peaks belonging to a single gene ontology term is shown in brackets beside the term. TSS, transcriptional start sites; GO, gene ontologies; DEGs, differentially expressed genes.

Histone modifications induced by environmental influences have been reported to be stable throughout somatic cell division (Abdelsamed et al., 2018; Almouzni and Cedar, 2016; Gensous et al., 2019; Pal and Tyler, 2010; Vickers, 2014). Hence, we investigated whether the effects of IF on H3K9me_3_ were maintained following refeeding. Interestingly, our findings revealed that mice subjected to refeeding (IF16.R and EOD.R) continue to display differential occupancy sites as compared to the control group (AL.R), suggesting that the modulation of gene expression by H3K9me_3_ was maintained after termination of IF (**Figure 2d**). The ChIP-seq data profiles at the H3K9me_3_ mark around TSSs of annotated genes were strikingly different for IF16.R and EOD.R as compared to AL.R (**Figure 2e**). Overall, histone marks were downregulated in IF16.R but upregulated in EOD.R (**Figure 2e**). Following refeeding, 219 differential peaks were identified for IF16.R and 47 differential peaks in EOD.R as compared to AL.R (**Figure 2f**). Of these, 7 differential peaks were common to IF16.R and EOD.R (**Figure 2f**). Functional analysis of genes associated with the differential peaks for IF16 and EOD showed enrichment of Gene Ontology (GO) terms relevant to metabolic processes and cellular transportation (**Figure 2g**), whereas terms related to redox processes, metabolic processes, DNA damage and repair response, and transcriptional-related events were enriched for the genes associated with the differential peaks for IF16.R and EOD.R (**Figure 2h**).

### 2.3 Transcriptomic analysis of IF and refeeding in the cerebellum

Following the changes detected in the epigenetic landscape, we next profiled the global cerebellum transcriptome by RNA-sequencing to ascertain the alterations in gene expression following IF and refeeding. Partial least square-discriminant analysis (PLS-DA) grouped the data into three clusters largely corresponding to the three study groups, AL, IF16 and EOD (**Figure 3a**). The clustering patterns suggested that IF16 and EOD induced differential effects on global gene expression as compared to AL. Notably, unsupervised hierarchical clustering of the data also resulted in distinct segregation into AL, IF16 and EOD groups (**Figure 3b**). Global gene expression profiles of IF16 and EOD were more similar to each other and strikingly different from that of AL, although specific clusters of genes showed stark differences between IF16 and EOD (**Figure 3b**), suggesting that IF induces time-dependent changes to the transcriptome. This finding is consistent with the results of the PLS-DA analysis and further reinforces the findings from the blood analyses. Volcano plots on the results of differential expression analysis (**Figure 3c**) identified 892 significantly differentially expressed genes in IF16 as compared to AL. Of these, 453 genes were up-regulated and 439 genes were down-regulated in IF16 (**Figure 3c**). On the other hand, 1472 genes were significantly differentially expressed in EOD as compared to AL; of these, 552 genes were up-regulated and 920 genes were down-regulated in EOD (**Figure 3c**). We carried out the GO enrichment analysis for the differentially expressed genes to identify the biological processes affected by both IF regimens. The top 20 enriched GO terms for IF16-induced differentially expressed genes were associated with circadian rhythm process, transcription related events, and steroid lipid metabolic process (**Figure 3d**). In addition to transcription related events and steroid lipid metabolic process, the differentially expressed genes in EOD were associated with cell proliferation, and differentiation through modulation of signaling pathways such as Ras protein, platelet-derived growth factor receptor (PDGFR) and epidermal growth factor receptor (EGFR) signaling pathways, suggesting a wider repertoire of transcriptional modulation with prolonged IF (**Figure 3d**). In summary, IF modulates the transcriptome in a time-dependent manner in agreement with the results of the ChIP-seq analysis.

**Figure 3:**
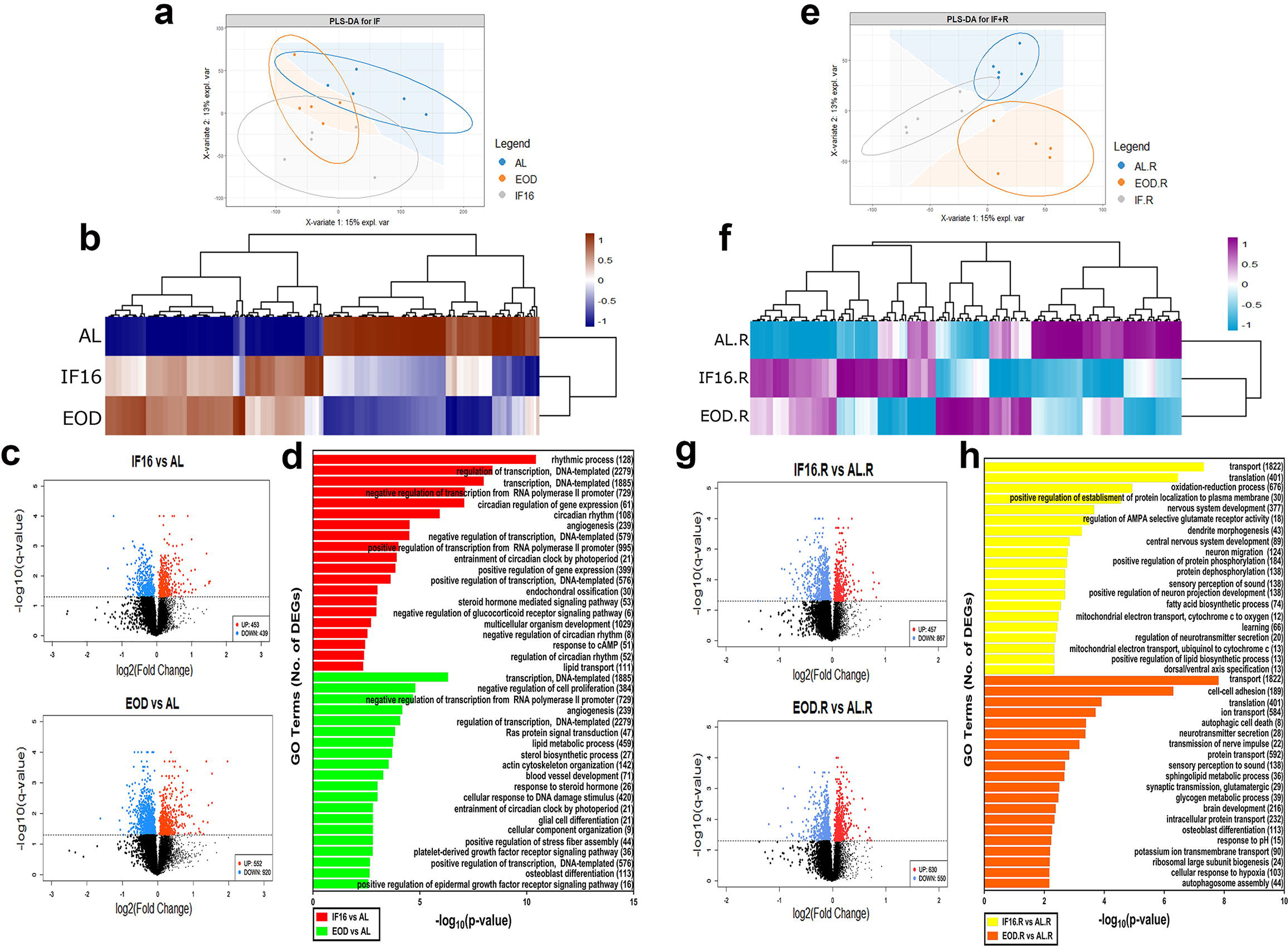
Transcriptomic analysis of IF and refeeding in the cerebellum. (a) Partial least square-discriminant analysis (PLS-DA) plot of IF transcriptomic expression profiles. Axis values are the explained variation of each variate. PLS-DA categorized AL, IF16, and EOD transcriptomic datasets into three unique cluster regions represented by the respective ellipse and background color. (b) Heatmap of transcriptomic expression data showing IF differentially expressed genes. Unsupervised hierarchical clustering segregated AL, IF16, and EOD transcriptome distinctly. Gene expression is shown in log_10_(FPKM+1) and differentially expressed genes were selected based on p-value<0.05. (c) Volcano plot of statistical significance (−log_10_ q-value) against enrichment (log_2_ fold change) of differentially expressed genes in IF16 and EOD against AL. Total number of differentially expressed genes shown in brackets. Upregulated genes are shown in orange and downregulated genes are shown in blue. Non-significant differentially expressed genes are shown in black. (d) Top 20 differentially expressed gene ontologies of IF compared to control plotted against statistical significance (represented as (−log_10_ p-value). Number of differentially expressed genes belonging to a single gene ontology term is shown in brackets beside the term. (e) PLS-DA plot of refeeding transcriptomic expression profiles. Axis values show the explained difference between each variate. PLS-DA categorized AL.R, IF16.R, and EOD.R transcriptomic datasets into three unique cluster regions represented by the respective ellipse and background color. (f) Heatmap of transcriptomic expression data showing refeeding differentially expressed genes. Unsupervised hierarchical clustering distinctly segregated AL.R, IF16.R and EOD.R transcriptomes. Gene expression is shown in log_10_(FPKM+1) and differentially expressed genes were selected based on p-value<0.05. (g) Volcano plot of statistical significance (−log_10_ q-value) against enrichment (log_2_ fold change) of differentially expressed genes in IF16.R and EOD.R against AL.R. Total number of differentially expressed genes shown in brackets. Upregulated genes are presented in red and downregulated genes are presented in blue. Non-significant differentially expressed genes are shown in black. (h) Top 20 differentially expressed gene ontologies of refeeding compared to control plotted against statistical significance (represented as (−log_10_ p-value). Number of differentially expressed genes belonging to a single gene ontology term is shown in brackets beside the term. FPKM, fragments per kilobase of transcript per million mapped reads.

Fasting and refeeding has been reported to have differential effects on the transcriptome profile in different organisms and organs (Kinouchi et al., 2018; Yang et al., 2019; Zhang et al., 2011). We examined the impact of refeeding on the transcriptome and notably found using PLS-DA analysis distinct clusters of AL.R, IF16.R and EOD.R with minimal overlap, indicating significant differences in the transcriptome profiles across the groups in spite of similar and abundant food intake (**Figure 3e**). Unsupervised hierarchical clustering further showed distinct expression profiles across the three groups, (**Figure 3f**). Volcano plots identified 1326 significantly differentially expressed genes between IF16.R and AL.R; of these, 457 genes were up-regulated and 867 genes were down-regulated in IF16.R (**Figure 3g**). On the other hand, 1180 genes were significantly differentially expressed between EOD.R and AL.R; of these, 630 genes were up-regulated and 550 genes were down-regulated in EOD.R (**Figure 3g**). Functional analysis of the differentially expressed genes in IF16.R as compared to AL.R showed that they were associated with cellular transportation process, translation related events, nervous system development including dendrite morphogenesis and neuron migration and projection, neurotransmitter activities and learning as well as fatty acid biosynthesis and oxidative phosphorylation (**Figure 3h**). On the other hand, differentiation of genes propagated by EOD.R as compared to AL.R was related to neurotransmitter activities, cellular transportation process, translational-related events, sphingolipid and glycogen metabolic process, as well as autophagy mechanisms (**Figure 3h**). These results indicate that fasting and refeeding have heterogeneous effects on the transcriptome, and provide further evidence that these regimens induce differential adaptive molecular responses to energy restriction and abundance.

### 2.4 IF and refeeding regulate distinct as well as common biological processes

We carried out an integrative analysis of the ChIP-seq datasets from IF and refeeding mice to compare the epigenetic landscape at H3K9me_3_ sites following the two regimens. 3D-PCA plot shows a clear separation of the ChIP-seq dataset from each biological condition (**Figure 4a**). Notably, IF16 and IF16.R were closer to each other as compared to EOD and EOD.R. Results obtained from the normalized strand cross-correlation curve show strong fragment-length peak whereas fingerprint plot demonstrate patterns of broad repressive mark typical of H3K9me_3_, highlighting high quality control of ChIP-seq dataset **(Figure 4b and c)**. Moreover, normalized strand cross correlation curve demonstrated a higher focal enrichment signal for IF16 and a lower localized enrichment signal for EOD, as compared to AL (**Figure 4b**). Overall, the enrichment signal of the IF ChIP-seq dataset was apparently higher than that of the refeeding dataset, which was less distinct across the biological conditions (**Figure 4c**). Normalized heatmap analysis across the groups showed that the overall histone marks patterns around the H3K9me_3_ locus were enriched in IF16, but downregulated in EOD, relative to AL (**Figure 4d**). In contrast, the overall gene expression pattern for IF16.R was downregulated while EOD.R was upregulated, as compared to AL.R (**Figure 4d**). Further, global gene expression around the TSS at H3K9me_3_ locus was generally higher for refeeding groups than IF groups. Broadly, these results further suggest that the regimens exert a differential impact on the H3K9me_3_ landscape.

**Figure 4:**
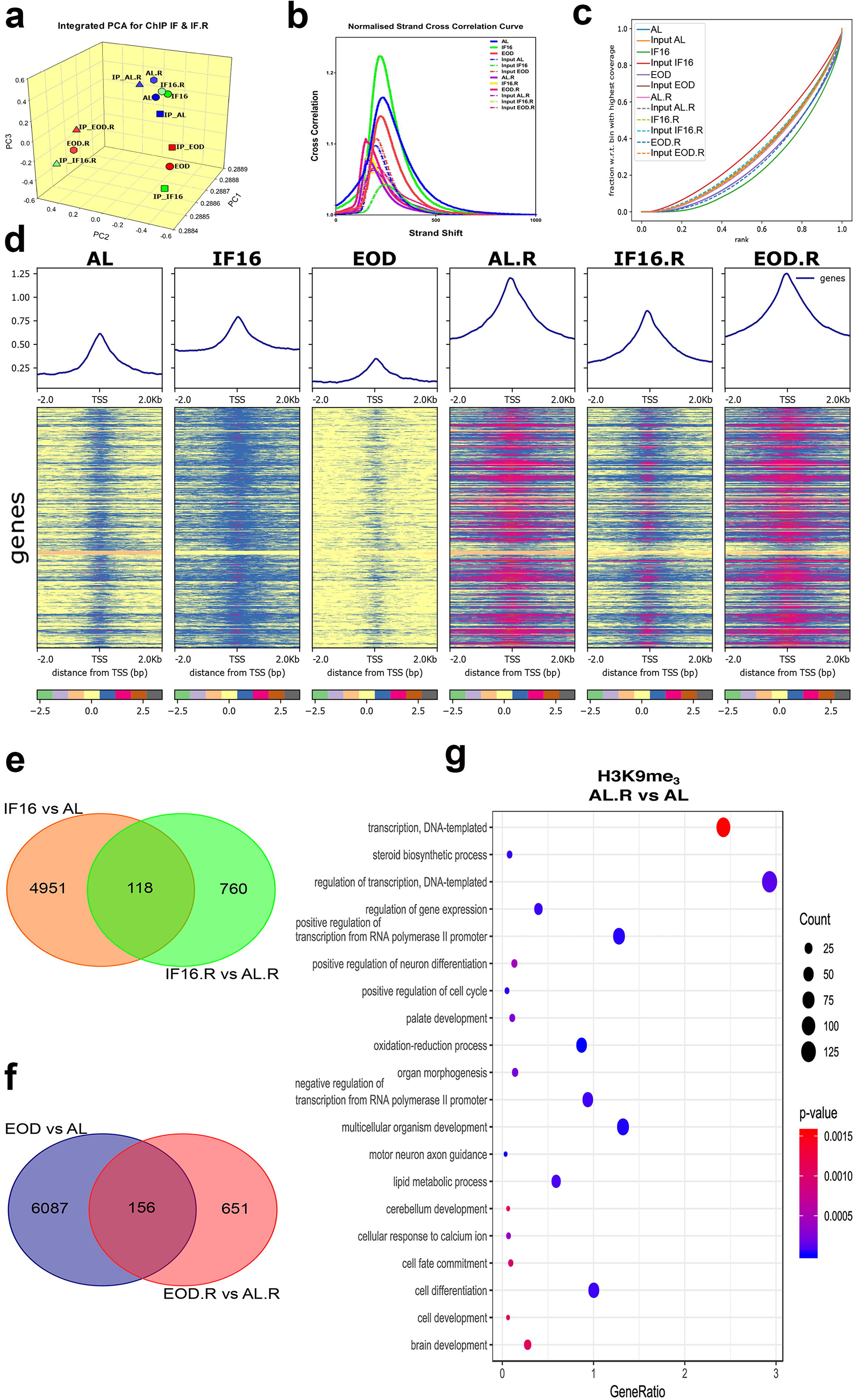
Integrative epigenomic analysis at the H3K9me_3_ locus in the cerebellum, on IF and refeeding. (a) 3D-PCA plot of normalized H3K9me_3_ ChIP dataset shown by expression profiles of IF and refeeding groups. The three most significant principal components (PC1, PC2, and PC3) are displayed on the x-, y- and z-axes, respectively. PCA categorized all biological groups through unique cluster occupancy relative to their input control. (b) Normalized strand cross correlation (SCC) plot for ChIP-seq samples of IF and refeeding. The ChIP-seq SCC curves show local maxima in the fragment sizes. (c) Fingerprint plots assess the relative focal signal strength and specificity of ChIP signal vs input signal of both IF and refeeding dataset. Both input and biological group datasets demonstrate good coverage of reads across the genome. (d) Summary and heatmap plots displaying normalized H3K9me_3_ ChIP-seq signal mapping to a 2 kb window around the TSS of genes revealed distinct expression pattern in biological samples of IF and refeeding groups. Normalised ChIP-seq signal over the body of genes are sorted according to mean score and scale bar illustrate log_2_ ratio of ChIP signal vs control signal. (e-f) Venn diagram illustrates the number of differentially expressed peaks at the H3K9me_3_ locus that are common and distinct between IF and refeeding regimen. (g) Dot plot for age-associated changes on differentially-expressed peaks ontologies. Gene ontologies semantic is shown on the left axis whereas Gene ratio is show on the x-axis. Dot size is proportional to the number of genes, and colour is presented as p-value.

We carried out a network analysis on the functional annotations of genes associated with differential peaks (**Figure S4**). Cluster profiling on the resulting network revealed that genes associated with differential peaks in IF16, as compared to AL, orchestrate a myriad of functions (**Figure S4a**). These functions can be categorized into several metabolic processes, such as fatty acid, glycerolipid, carbohydrate (glucose and glycogen), and protein metabolism. The genes associated with the differential peaks were also related to a plethora of signaling pathways such as G-protein coupled receptor (GPCR), Janus kinase (JAK) and signal transducer and activator of transcription protein (STAT), bone morphogenetic protein (BMP), Wnt, and integrin-mediated signaling pathways. Furthermore, they also facilitate transcription-related events, transport, circadian rhythm and chromatin organization. Cluster profiling of genes associated with the differential peaks in IF16.R, as compared to AL.R, showed fewer functional annotations (**Figure S4a**). These functional annotations showed terms associated with metabolic processes such as glycogen, peptide, quinolinate, collagen and chondroitin sulfate proteoglycan metabolism. Other functions of the differential peak-associated genes included those related to the GPCR signalling pathway, cyclic adenosine monophosphate (cAMP) kinase activity, pyrimidine nucleotide transport into mitochondrion, as well as small RNA interference. Despite fewer clusters being present in the profiling output following refeeding, we observed a total of 118 differential peaks that were maintained following refeeding after the IF16 regimen (**Figure 4e**). Cluster profiling of these differential peaks highlighted functions related to lipoprotein lipase activity, protein ubiquination, quinolinate catabolic process, tetrahydrofolate metabolism, and glycogen biosynthetic process (**Figure S4a**).

Compared to AL, EOD differential peaks were associated with functions related to lipid metabolism (e.g. fatty acid biosynthetic process, low-density lipoprotein particle receptor catabolic process, and sterol metabolic process and signalling), carbohydrate metabolism (e.g. malate and acetyl-coA metabolic processes), protein catabolism, tetrahydrofolate metabolic process as well as mucopolysaccharide and heparan sulfate proteoglycan metabolic processes (**Figure S4b**). These differential peaks were also associated with neurotransmitter biosynthesis (e.g. serotonin) and endosomal transport, complement activation, histone modification and RNA processing, and regulation of the Wnt signaling pathway. Furthermore, cluster profiling of EOD.R differential peaks as compared to AL.R, revealed fewer functional annotations compared to EOD vs. AL (**Figure S4b**). These differential peaks were enriched for glycogen and proteoglycan biosynthetic process, and quinolinate and tetrahydrofolate metabolic processes. Moreover, these differential peaks were related to the GPCR signalling pathway, DNA repair, anatomical structure development and sensory perception of mechanical stimuli (**Figure S4b**). A total of 156 differential peaks were maintained between EOD and EOD.R, when compared to AL and AL.R, respectively (**Figure 4f**). These peaks mediated key metabolic processes, such as carbohydrate metabolism (e.g. carbohydrate utilization, and GDP L-fucose and glycogen metabolic processes), quinolinate and exogeneous drug catabolism as well as catalyzing processes, such as equilibrioception, RNA interference, complement activation and neurotransmitter secretion **(Figure S4b)**.

To determine whether age-associated changes may have a confounding effect on the observed functional annotations, we investigated age-associated differential peaks between AL and AL.R (**Figure 4g**). Our results revealed significant ontologies belonging to terms such as lipid and steroid metabolic process, transcriptional-related events, organism development processes (e.g. palate, organ morphogenesis, motor neuron axon guidance, brain and cerebellum development) as well as redox processes. Many of these age-associated changes were evident in functional ontologies of both IF16.R and EOD.R, as well as in IF. Our findings thus raise the possibility that age is an important factor influencing epigenetic reprogramming at the H3K9me_3_ locus, and it may have a confounding effect on the changes brought about by refeeding. Thus, it is unlikely that the epigenetic maintenance reported in this study is due only to IF without any age-associated effects.

### 2.5 Integrative transcriptomic dataset of IF and refeeding demonstrate distinct and common regulation of a plethora of metabolic processes

We next conducted an integrative transcriptomic analysis between the IF and refeeding datasets. The PLS-DA plot showed that the AL, IF16, and EOD transcriptomic datasets were separated from each other, occupying three unique cluster regions (**Figure 5a**). Heat map analysis revealed a stark variation in gene expression patterns among the different groups (**Figure 5b**). Gene network analysis revealed that genes differentially expressed in IF16 compared to AL, belonged to various functional clusters spanning metabolic processes, such as fatty acid catabolism, sphingolipid metabolism, amino acid and mitochondrial pyruvate transport, as well as protein deubiquitination and ubiqutination (**Figure S5a**). In IF16.R, genes differentially expressed compared to AL.R were related to protein ubiquitination and peptide and ATP metabolism (**Figure S5a**), among other processes. A total of 83 differentially expressed genes were maintained between IF16 and IF16.R when compared to AL and AL.R respectively (**Figure 5c**). These genes were involved in a plethora of metabolic processes, such as 3’-phosphoadenosine 5’-phosphosulfate and S-adenosylmethionine biosynthesis, quinolinate catabolism, as well as xylulose and estrogen metabolic processes (**Figure S5a**). In addition, genes involved in GPCR and Hippo signalling pathways, transport (e.g. zinc ion import into synaptic vesicles, nuclear retention of unspliced RNA and urea transmembrane transport), cellular differentiation (e.g. B cell and epithelial cell differentiation), replication fork maintenance, interferon-gamma secretion, cellular response to chromate, and collagen fibril organization also showed maintenance between the feeding conditions.

**Figure 5:**
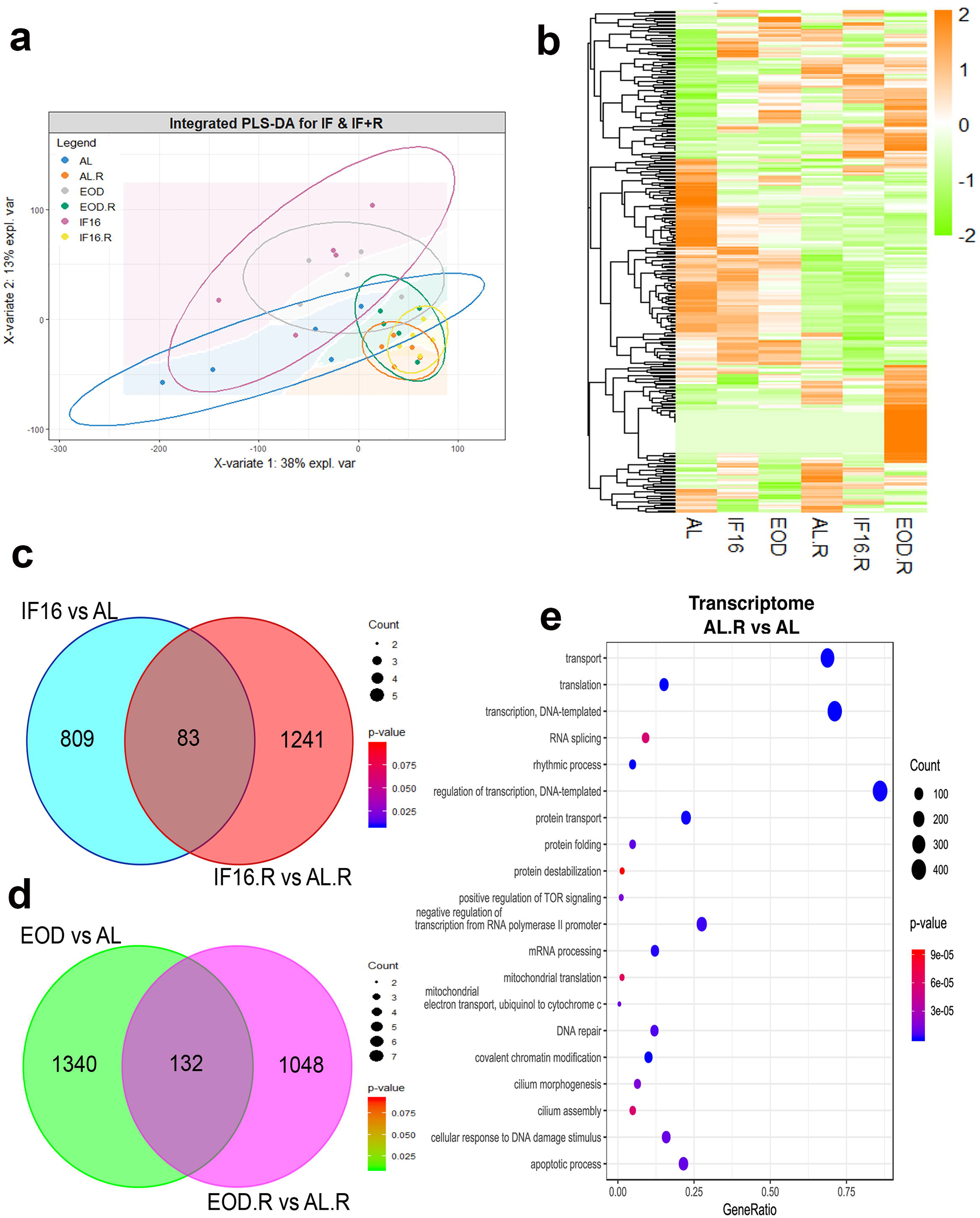
Integrative transcriptomic analysis of IF and refeeding in the cerebellum. (a) PLS-DA plot of normalised IF and refeeding transcriptomic expression profiles. Axis values are the explained differences in each variate. PLS-DA categorized AL, IF16, and EOD datasets, distinctly represented by the respective ellipse and background color; yet AL.R, IF16.R, and EOD.R transcriptomic dataset demonstrates little segregation. (b) Heatmap of normalized IF and refeeding transcriptomic expression data showing differentially expressed genes across biological groups. Gene expression is shown in log_10_(FPKM+1) and differentially expressed genes were selected based on p-value<0.05. (c-d) Venn diagram illustrates the number of differentially expressed genes that are common and distinct between IF and refeeding regimens. (e) Dot plot for age-associated changes on differentially-expressed gene ontologies. Gene ontologies semantic is shown on the left axis whereas Gene ratio is shown on the x-axis. Dot size is proportional to the number of genes, and colour is presented as p-value.

Compared to AL, EOD-induced genes differentially expressed were involved in steroid biosynthesis, protein ubiquitination, and glycosylphosphatidylinositol metabolism (**Figure S5b**). These genes also mediated GPCR and TOR signaling pathways, anatomical structure development, as well as ribosome biogenesis. Conversely, relative to AL.R, EOD.-induced differentially expressed genes were involved in metabolism-related activities, such as peptide biosynthesis and transport, protein ubiquination, RNA biosynthesis, as well as generation of precursor metabolites and energy, and GPCR signaling pathways (**Figure S5b**). Despite a lower gene ontology output compared to both IF16 and IF16.R, a greater number of differentially expressed genes was maintained (a total of 132) between EOD and EOD.R (**Figure 5d**). Notably, these genes were involved in metabolism (e.g. citrate metabolic process, amylopectin biosynthesis, quinolinate catabolism, protein ubiquination, dipeptide and lipid transport), transcription-related events, and the GPCR and target of rapamycin complex 1 (TORC1) signaling pathways (**Figure S5b**). These genes also mediate negative regulation of other processes (e.g. dendritic spine maintenance, FasL and CD4 biosynthesis), modifications (e.g. histone succinylation, tRNA aminoacylation for protein translation, peptidyl-aspartic acid hydroxylation and peptidyl-glutamic acid modification), anatomical structure development, regulation of muscle system process, cerebellar granule cell precursor proliferation, cellular response to glial cell derived neurotrophic factor, sodium transport, and establishment of cell polarity.

Since we observed age-associated changes the epigenome, we investigated if age is a confounder for the transcriptome dataset (**Figure 5e**). Gene ontologies of age-associated differentially expressed genes belonged to transcription and translation events, transportation, DNA damage and repair responses, mitochondrial electron transport, TOR signalling, protein folding and destabilization, and apoptosis. Many of these gene ontologies overlapped with some of the terms enriched during refeeding and maintained differentially expressed genes suggesting a possible influence of age in addition to feeding effects.

### 2.6 Integrative epigenomic and transcriptomic dataset highlight robust metabolic processes regulated by H3K9me_3_ modulation during IF in a temporal-dependent manner

We next investigated the profile of differentially expressed genes in both regimens that were regulated by H3K9me_3_ modulation using integrative multi-omics. A total of 208 differentially expressed genes governed by H3K9me_3_ were found to be common between IF16 and AL (**Figure 6a**). As IF modulation of the H3K9me_3_ mark resulted in robust metabolic process changes, we focused on gene ontologies related to the term “metabolism” and categorized them into three major arms: carbohydrate, lipid, and protein (**Figure 6b**). Of the 208 differentially expressed genes, 89 were responsible for regulating metabolism. Between IF16 and AL, genes associated with carbohydrate and steroid metabolic processes were upregulated, whereas those involved in lipid and protein metabolic processes were downregulated (**Figure 6b**). Also, many of these genes were involved in more than one function, suggesting that the conglomerate of these 89 genes is necessary to reflect the overall metabolic changes observed. We then broadly separated both the upregulated and downregulated differentially expressed genes governed by H3K9me_3_ locus and analyzed their pathway changes related to metabolism (**Figure 6c**). Notably, upregulated genes mediated pathways related to circadian rhythm, various signalling arms (e.g. mitogen-activated protein kinases [MAPK], Ras-related protein 1 [Rap1] and Ras), as well as aspects of metabolism (e.g. sphingolipid, sulfur and vitamin B6). Conversely, downregulated genes were involved in sphingolipid, phospholipase D, mTOR, Hippo and adipocytokine pathways, as well as the regulation of lipolysis in adipocytes and bile secretion (**Figure S6**). Only 25 H3K9me_3_-controlled differentially expressed genes were observed between IF16.R and AL.R (**Figure 6d**). However, gene ontologies of these 25 genes revealed only two broad categories of positive regulation of the apoptotic signaling pathway and regulation of DNA-templated transcription (**Figure 6e**). Overlap analysis of these H3K9me_3_ regulated differentially expressed genes between IF16 and IF16.R, relative to AL and AL.R, respectively, investigating if metabolic changes were maintained following refeeding showed that no differentially expressed genes intersected between the two regimens (**Figure 6f**).

**Figure 6:**
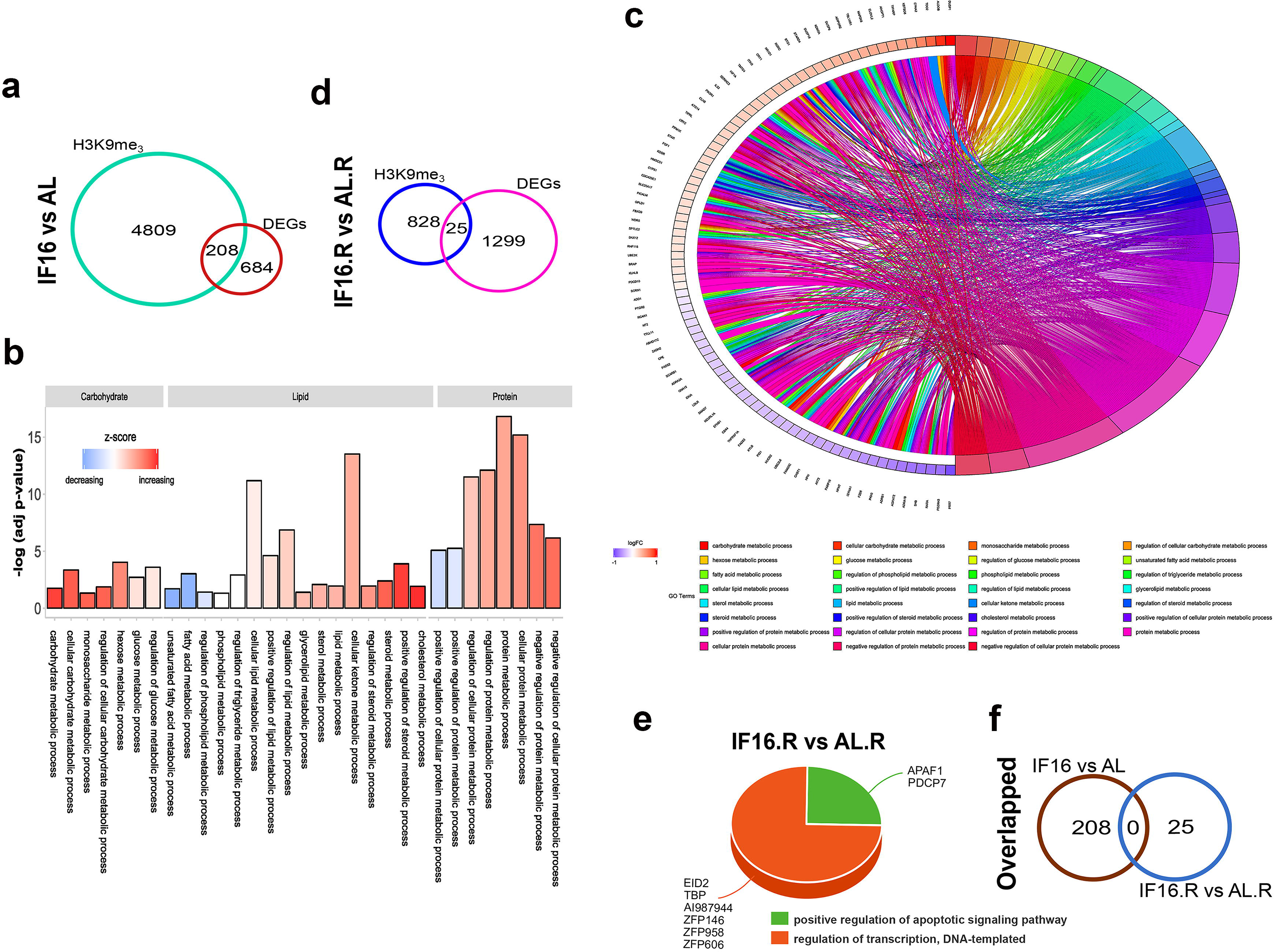
Integrative epigenomic and transcriptomic analysis of IF16 and IF16.R at the H3K9me_3_ locus in the cerebellum. (a) Euler’s diagram illustrates the number of differentially expressed genes governed by H3K9me_3_ modulation during IF16 compared to control. (b) Bar chart categorising H3K9me_3_-governed differentially expressed genes during IF16 vs AL into three categories of metabolic gene ontologies; namely carbohydrate, lipid and protein metabolic processes. Gene ontology subsets of each category are shown at the bottom of each bar chart. Statistical significance is plotted as −log_10_(adj p-value) on the y-axis whereas z-score is used in the form of a colorbar to illustrate whether a particular semantic term is more likely to increase or decrease. Red represents an increasing z-score whereas blue represents a decreasing z-score. (c) Chord diagram showing the most enriched carbohydrate, lipid and protein related metabolic processes with H3K9me_3_-governed differentially expressed genes during IF16 vs AL. In each chord, enriched gene ontologies are presented on the right, whereas differentially expressed genes contributing to this enrichment are presented on the left. Each differentially expressed gene is represented by a rectangle in which the color is correlated to the expression level determined by log(fold-change), in which red represents upregulation and blue represents downregulation. Chords connect these differentially expressed genes with gene ontology terms, with each term being represented by one coloured line. (d) Euler’s diagram illustrates the number of differentially expressed genes governed by H3K9me_3_ modulation during IF16.R compared to control. (e) 3D-pie chart to illustrate H3K9me_3_-governed differentially expressed genes during IF16.R vs AL.R. Gene ontologies are presented at the bottom of the pie chart, whereas gene symbols belonging to each semantic gene ontology are shown at each category of the pie chart. (f) Venn diagram illustrates the number of differentially expressed genes that are governed by H3K9me_3_ modulation that are maintained between IF16 and IF16.R compared to that in control.

Similar pipeline analysis for EOD and EOD.R revealed 423 differentially expressed genes to be governed by H3K9me_3_ between EOD and AL (**Figure 7a**). Gene ontology analysis revealed upregulation of carbohydrate, protein, fatty acid and triglyceride metabolic processes, and downregulation of steroid metabolism (**Figure 7b**). Chord diagram analysis revealed that 175 of the 423 differentially expressed genes were responsible for regulating metabolism (**Figure 7c**). Two other notable details were observed in the case of EOD, compared to IF16. EOD showed a large number of differentially expressed genes, involved in a number of metabolic processes; many of the upregulated genes controlled the protein metabolic axis (**Figure 7c**). Analysis of the differentially expressed genes revealed more upregulated terms than downregulated terms. Upregulated differentially expressed genes governed a plethora of signaling pathways (e.g. Rap1, Forkhead box O [foxO], sphingolipid, longevity regulating, cyclic adenosine monophosphate [cAMP], 5’-adenosine monophosphate kinase [AMPK], Ras, MAPK, phosphatidylinositol, Toll-like receptor, glucagon, mTOR, thyroid hormone, adipocytokine, and relaxin), as well as circadian rhythm, sphingolipid metabolism, folate and unsaturated fatty acids biosynthesis, insulin resistance, parathyroid hormone synthesis, secretion and action, and fatty acid elongation. Downregulated differentially expressed genes mediated pathways related to steroid and unsaturated fatty acids biosynthesis, metabolism (e.g. cysteine, methionine, pyrimidine, glutathione), vitamin digestion and absorption, as well as circadian rhythm (**Figure S6b**). H3K9me_3_-controlled differentially expressed genes analysis revealed 18 genes between EOD.R and AL.R (**Figure 7d**), categorized into two broad semantics: phospholipid metabolic process and myelination (**Figure 7e**). An overlap analysis of H3K9me_3_-regulated differentially expressed genes between EOD and EOD.R, relative to AL and AL.R respectively, found no differentially expressed genes that intersected between the two regimen (**Figure 7f**).

**Figure 7:**
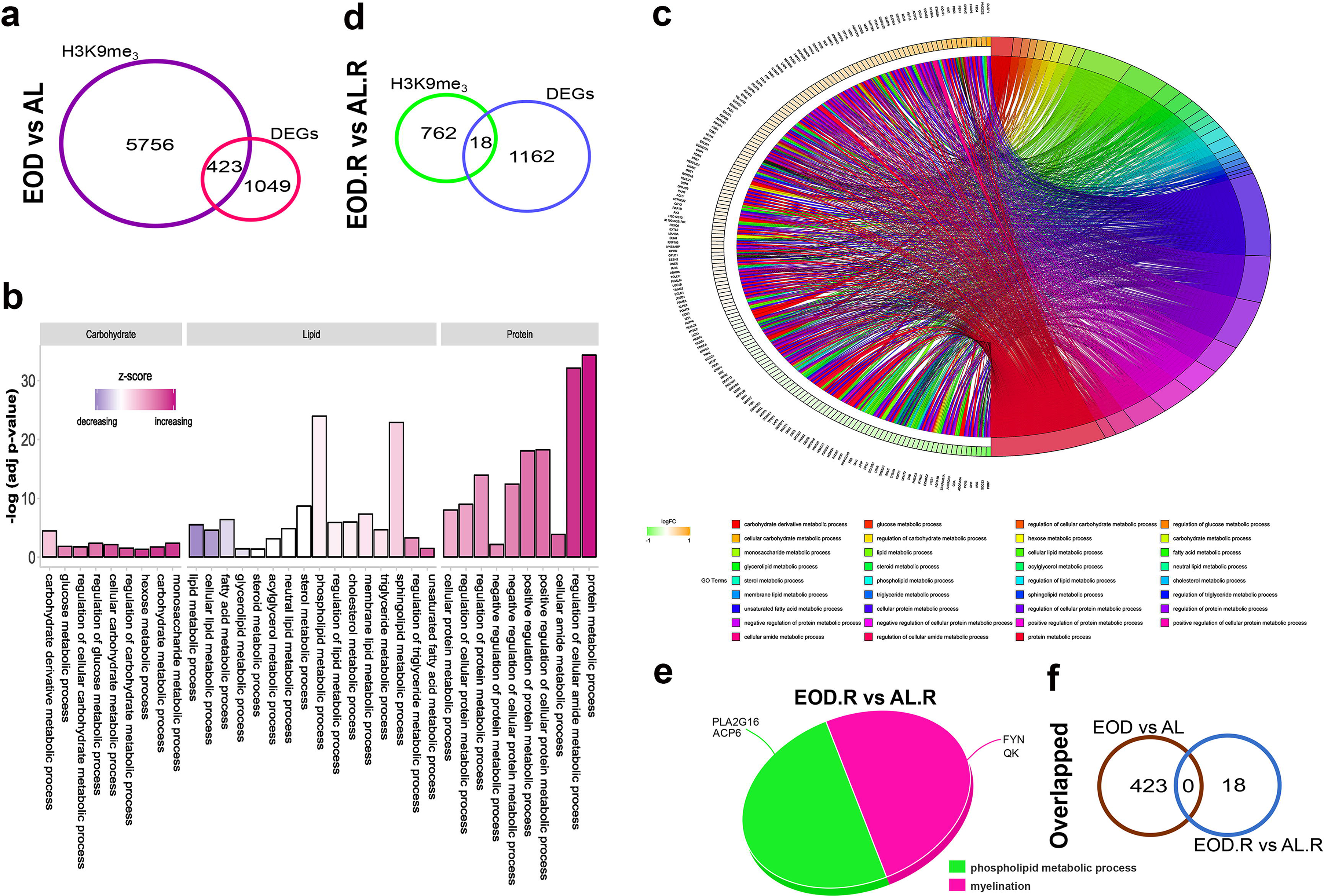
Integrative epigenomic and transcriptomic analysis of EOD and EOD.R at the H3K9me_3_ locus in the cerebellum. (a) Euler’s diagram illustrates the number of differentially expressed genes governed by H3K9me_3_ modulation during EOD compared to control. (b) Bar chart categorising H3K9me_3_-governed differentially expressed genes during EOD vs AL into three categories of metabolic gene ontologies; namely carbohydrate, lipid and protein metabolic processes. Gene ontologies subset of each category are shown at the bottom of each bar chart. Statistical significance is plotted as −log_10_(adj p-value) on the y-axis whereas z-score is used in the form of a colorbar to illustrate whether a particular semantic term is more likely to increase or decrease. Purple represents increasing z-score whereas cyan represents a decreasing z-score. (c) Chord diagram showing the most enriched carbohydrate, lipid, and protein-related metabolic processes with H3K9me_3_-governed differentially expressed genes during EOD vs AL. In each chord, enriched gene ontologies are presented on the right, whereas differentially expressed genes contributing to this enrichment are presented on the left. Each differentially expressed gene is represented by a rectangle whose color is correlated to the expression level determined by log(fold-change), in which red represents upregulation whereas blue represent downregulation. Chords connect these differentially expressed genes with gene ontology terms, with each term being represented by one coloured line. (d) Euler’s diagram illustrates the number of differentially expressed genes governed by H3K9me_3_ modulation during EOD and EOD.R compared to control. (e) 3D-pie chart to illustrate H3K9me_3_-governed differentially expressed genes during EOD.R vs AL.R. Gene ontologies are presented at the bottom of the pie chart, whereas gene symbols belonging to each semantic gene ontology are shown at each category of the pie chart. (f) Venn diagram illustrates the number of differentially expressed genes that are governed by H3K9me_3_ modulation that are maintained between EOD and EOD.R compared to that in control.

As there was a temporal difference in the modulation of differentially-expressed genes by H3K9me_3_ in both IF16 and EOD, to reinforce our findings we looked at plausible interactions between H3K9me_3_ peaks with the transcriptome track of selected genes (**Figure 8**). Evaluation of representative genes distinctly modulated only in IF16 and EOD (e.g. *Aldob* and *Gbe1* respectively), showed that a peak change at the H3K9me_3_ locus resulted in upregulated expression of both genes (**Figure 8a-b**). Next, we compared representative genes that were modulated in both IF16 and EOD (e.g. *Elovl2* and *Fads2*) as a result of H3K9me_3_ modulation. Similar to previous observations, H3K9me_3_ modulation resulted in upregulation of *Elovl2* expression and downregulation of *Fads2* expression (**Figure 8c-d**), highlighting the distinct impact of varying the period of energy restriction on relative gene expression patterns. Thus, IF can affect gene expression in a temporal manner impacting metabolic changes differently. Indeed, when we mapped the differentially expressed genes from both IF regimens that were governed by H3K9me_3_ changes onto a metabolic map, more nodes were present during EOD than IF16, and the metabolic profile was very different (**Figure S7a-b**). During IF16, most differentially expressed genes were involved in carbohydrate metabolism, whereas in EOD there was a metabolic switch from carbohydrate metabolism to lipid and amino acid metabolism (**Figure S7b**). When we focused on the anabolic and catabolic changes of these metabolic pathways as a whole, we observed that the metabolic utilization of key macromolecules differed greatly between IF16 and EOD. For instance, for EOD, biosynthesis of carbohydrate, amine, and polyamine was greater than for IF16, but biosynthesis of amino acids, fatty acids, and lipids was less (**Figure S8a**). On the other hand, EOD degraded less carbohydrate, amines, and polyamines than IF16, but degraded more amino acids, fatty acids, and lipids than IF16 (**Figure S8b**). Collectively, there was a loss of maintenance of differentially expressed genes at the H3K9me_3_ locus following refeeding. This reinforces the notion that neither IF16 nor EOD are robust enough to induce epigenetic reprogramming at this locus to maintain the transcriptomic changes, even with the abolition of IF. Instead, it appears that IF16 and EOD induce metabolic changes to respond to energy restriction through modulation of the H3K9me_3_ locus in the cerebellum. However, our findings indicate that EOD can induce more robust metabolic switching than IF16, providing a novel insight that temporal regulation of H3K9me_3_ leads to distinct metabolic switching processes in the cerebellum.

**Figure 8:**
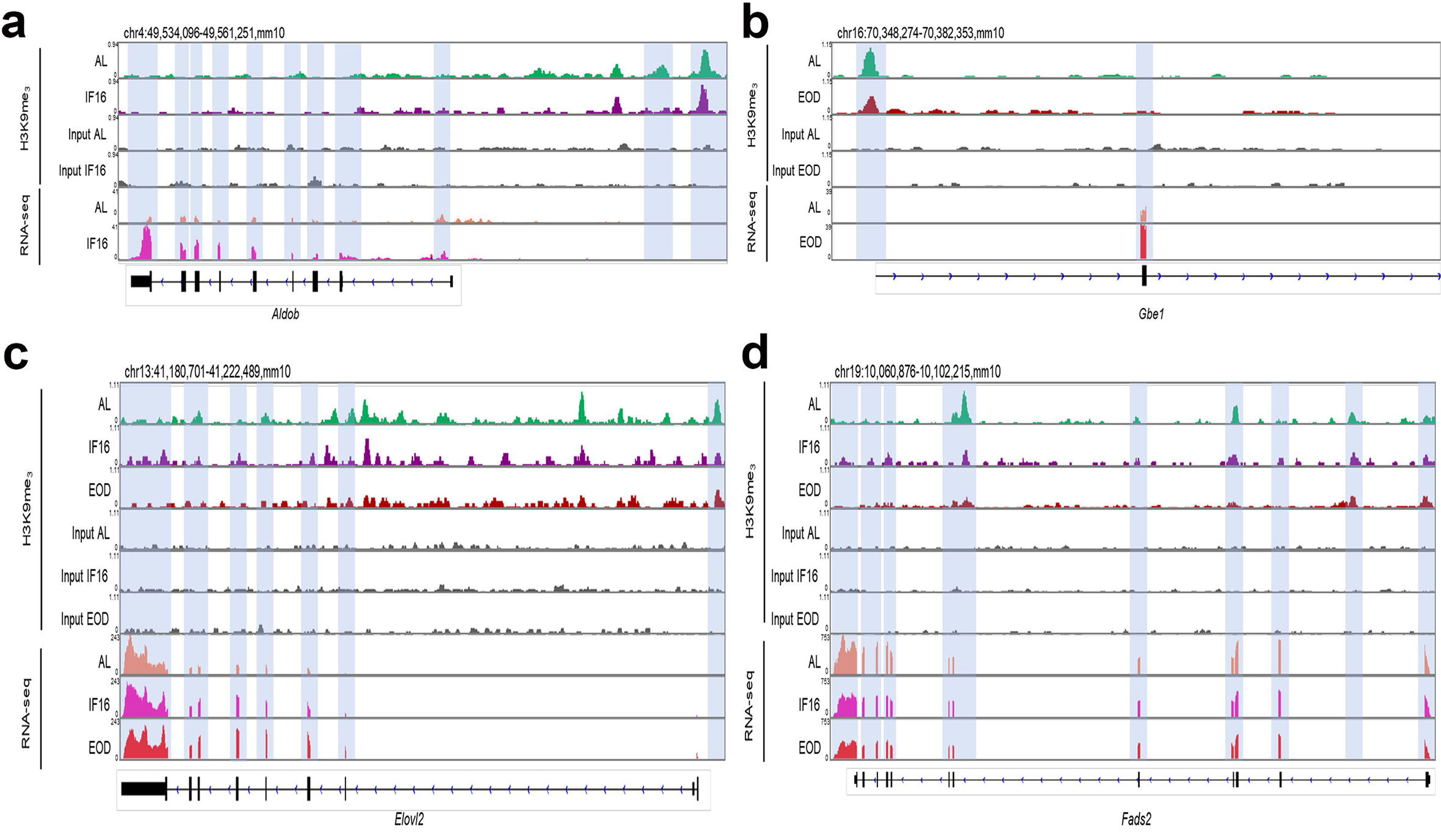
Integrative Genome Visualisation (IGV) tracks illustrating H3K9me_3_ binding sites at selected gene locus following IF. (a-b) IGV track displaying H3K9me_3_ binding sites at *Aldob* and *Gbe1* gene locus following IF16 and EOD respectively. (c-d) IGV track displaying H3K9me_3_ binding sites at *Elovl2* and *Fads2* gene locus following both IF16 and EOD. Blue background show H3K9me_3_-dependent regulation of chromatin accessibility at the selected sites. Location of selected gene locus is shown on the top.

## 3. Discussion

Accumulating evidence indicates that IF has numerous benefits for metabolic health, and is a plausible medical intervention. However, many studies may show different benefits of IF often because of confounding factors arising from differing periods of time-restricted feeding as well as inter-individual differences in response (Gabel et al., 2018; Journal et al., 2015; Paoli et al., 2019; Trepanowski and Bloomer, 2010). As such, the present study evaluated the temporal effects of IF by adopting different timepoints during IF and refeeding, and considered the epigenetic milieu of IF and refeeding to understand how this regulatory cascade may help to explain any response differences.

Considerable evidence has shown that CR and epigenetics are interlinked, providing a clearer understanding on various aspects of regulation of gene expression patterns through this axis. H3K9me_3_ is a key player in CR-induced modulation, which in turn affects metabolic adaptation and aging processes (Hernández-Saavedra et al., 2019; Maegawa et al., 2017; Molina-Serrano and Kirmizis, 2017). This epigenetic locus has also been widely implicated in metabolic modulation in different organs and disease states, suggesting that H3K9me_3_ has an important role in metabolic adaption (Strauss and Reyes-dominguez, 2014; Villeneuve et al., 2008; Yu et al., 2012). However, there is no available information on whether IF may also affect the epigenome. Here, we established that both IF and refeeding regimens can distinctly modulate the H3K9me_3_ tag. Our data indicate a novel mechanism in which both IF and refeeding influence metaboloepigenetics at the H3K9me_3_ locus, similar to observations in CR studies. Moreover, studies have reported that H3K9me_3_ can affect biological processes such as redox and DNA damage and repair response under excessive energy balance state (Gordon et al., 2015; 2018; Zhong et al., 2018). This was also observed in our study following refeeding. Our dataset is therefore consistent with other studies and indicates a role for H3K9me_3_ modulation as a notable player in this dietary context.

Recent studies have reported that the cerebellum affects circadian oscillation, which controls food anticipation behaviour (Delezie et al., 2016; Mendoza et al., 2010), and functions as an epigenetic clock during aging (Horvath et al., 2015). Given the large population of neuronal cells residing within the cerebellum and the corresponding energy demand, it is hypothesized that the cerebellum should be highly sensitive to energy restriction (Howarth et al., 2009, 2012; Kuzawa et al., 2014). Indeed, the transcriptomic dataset for both IF and refeeding, reveals robust changes in gene expression compared to control, indicating that both regimens act via a transcriptomic axis. It was also observed that the duration of fasting and refeeding distinctly affected the transcriptome. IF16-induced differentiation of genes was associated with the circadian rhythm process, which may explain the chronobiological changes in the cerebellum during food anticipation towards energy deprivation (Delezie et al., 2016; Mendoza et al., 2010). Moreover, both IF regimens modulated steroid lipid metabolic changes, highlighting metabolic switching processes typical of fasting, which in turn may affect neurodevelopment and neurodegeneration in the cerebellum (Kawamoto, 2016; Zhang et al., 2011). In addition, it was reported that 24-hour fasting induced stem cell regeneration by metabolic switching (Mihaylova et al., 2018). Here, we also established that EOD differentially expressed genes modulate cellular proliferation and differentiation by modulating signaling pathways such as Ras protein, PDGFR, and EGFR. Our data offer a transcriptomic lens to understanding the mechanistic aspects of how fasting impacts different signalling arms to regulate proliferation and differentiation in the cerebellum.

Integrative epigenetic and transcriptomic analyses of both fasting and refeeding datasets revealed both to have a profound effect on H3K9me_3_, but IF appears to have a greater effect on the transcriptome than refeeding. Network construction analysis of the epigenome shows that both IF16 and EOD induce H3K9me_3_ modulation, which in turn regulates fatty acid (e.g. fatty acid oxidation and biosynthesis), carbohydrate (e.g. acetyl-CoA and tricarboxylic acid cycle) as well as protein (e.g. protein catabolism) metabolism. At the transcriptomic level we also observe such changes in lipid (e.g. fatty acid catabolism and steroid biosynthetic process), carbohydrate (e.g. amino acid and mitochondrial pyruvate transport) and protein (e.g. protein deubiquination and ubiquination) metabolism suggestive of coordinated control of such metabolic events. Interestingly, our findings also show that, despite only an 8-hour difference in energy restriction, different facets of metabolic processes are distinctly modulated in both IF16 and EOD. For instance, we observed glucose and glycogen metabolic changes during IF16, but malate and acetyl-coA metabolic changes were only obereved during EOD as a result of H3K9me_3_ modulation. Therefore, different arms of cellular metabolic pathways are being modulated distinctly in different IF regimens to provide different energy metabolite sources to meet energy demands. Besides metabolism, both IF16 and EOD can induce other differential biological changes at the epigenomic and transcriptomic level, such as various signalling pathways, anatomical structure development, and RNA splicing events.

Refeeding following IF appears to also affect both the epigenome and transcriptome, governing several distinct biological processes in a temporal-dependent manner. Contrary to fasting, fewer genes and peaks were impacted upon refeeding, indicating a reduced effect on the epigenome and transcriptome. Moreover, certain functional maintenance was apparent at both the H3K9me_3_ locus as well as gene expression, suggesting that epigenetic reprogramming of gene expression may occur as a result of IF. Our study highlighted age as a potential confounding factor in the precise mapping of effects of refeeding following IF, as observed through differences in body weight and expression profiles of AL and AL.R mice. Indeed, many previous studies have also emphasized that aging has a profound effect on the level of H3K9me_3_ mark and transcriptome patterns in the cerebellum (Dillman et al., 2017; Maleszewska et al., 2016; Sidler et al., 2017; Snigdha et al., 2016). Consistent with these reports, we found that many differences in genes and peaks observed in mediating biological effects in refeeding overlap with age-associated biological changes. Hence, it is possible that age affects epigenetic reprogramming at the H3K9me_3_ locus, which makes it indeterminate that most of the epigenetic maintenance reported in this study between both types of regimen is induced fully by IF alone.

Collectively, our study shows for the first time that fasting can affect H3K9me_3,_ which regulates the many biological changes relevant to IF, some of which have been reported in different organ settings. In addition, since both IF regimens affected the epigenome and transcriptome distinctly, our data provide a plausible explanation that a distinct modulation of both the epigenome and transcriptome axis may produce different outcomes, which in turn may help to explain the lack of standardization in reported IF effects due to different IF regimens being employed. Given that several metabolic processes at both epigenomic and transcriptomic levels were modulated because of fasting, whether the transcriptomic changes governing these metabolic process changes are governed by H3K9me_3_ modulation need to be studied. Our multi-omics analysis revealed that differentially expressed genes from both IF regimens governed by H3K9me_3_ were involved in a variety of metabolic processes. Between IF16 and AL, carbohydrate and steroid metabolic processes appear to be upregulated, whereas lipid and protein metabolic processes were downregulated. However, comparison of EOD and AL revealed a metabolic shift whereby carbohydrate, protein, fatty acid and triglyceride metabolic processes were upregulated, with a corresponding downregulation of steroid metabolism. These metabolic changes in both IF regimens appeared to be orchestrated by unique as well as common genes. For instance, both *Aldob* (responsible for glycolysis (Le et al., 2018; Peng et al., 2008)) and *Gbe1* (responsible for glycogen storage (Akman et al., 2011; Iijima et al., 2018)), were uniquely upregulated in IF16 and EOD, respectively. However, in both IF16 and EOD common genes, such as *Elovl2* (involved in de novo lipogenesis, lipid storage, and subsequent fat mass expansion (Pauter et al., 2014; Rennert et al., 2018)) and *Fads2* (involved in biosynthesis of polyunsaturated fatty acids (Huang et al., 2017; Reynolds et al., 2018)), were upregulated and downregulated, respectively, in a temporal manner. In addition, different signalling arms were modulated distinctly in IF16 and EOD. For example, common pathways, such as the adipocytokine, mTOR, and sphingolipid metabolic pathways, were downregulated in IF16 but upregulated in EOD. In contrast, unique signalling pathways, such as FoxO and AMPK, were distinctly modulated only in EOD. For certain signalling pathways, we observed that the different genes were modulated in the two regimens. For instance, in the MAPK pathway, *Angpt1* was involved only in IF16, whereas *Egfr* was involved only in EOD. Indeed, global metabolic mapping revealed different metabolic profiles in IF16 and EOD, with asymmetrical trigger of different milieu of carbohydrate, lipid, and protein cellular metabolic pathways, as well as performing varying degrees of anabolism and catabolism of these macromolecules. While most of the signalling pathways have been previously reported to be influenced by fasting (Ahima et al., 2006; Bujak et al., 2015; Choi et al., 2018; Dogan et al., 2011; Goldstein et al., 2016; Guo et al., 2016; Jensen et al., 2019; Kim, 2009; Li et al., 2019; Luo et al., 2018; Mahadik, 2012; Martins et al., 2016; Miyazaki et al., 2004; Morikawa et al., 2004; Nicklin et al., 2013; Pel and Lines, 2015; Rodrı et al., 2012; Tulsian et al., 2018; Wee and Wang, 2017; Wijngaarden et al., 2019; Yoon et al., 2018), our findings suggest a temporal link of the signaling response to the period of energy restriction. This is achieved by manipulating the epigenetic and transcriptomic cascade differently, which in turn results in varied metabolic arms being triggered to respond to energy demand.

In the case of refeeding, differentially expressed genes governed by H3K9me_3_ were involved in positive regulation of apoptotic signalling and transcription in IF16.R, and phospholipid metabolism and myelination in EOD.R. Many studies have reported the role of H3K9me_3_ in influencing these biological processes (Black and Whetstine, 2011; Liu et al., 2015, 2017; Lu et al., 2018; Olcina et al., 2015; Ye et al., 2017). However, despite subjecting biological groups to excess energy balance following IF, it is observed that differentiation of certain genes is modulated distinctly in a temporal-dependent manner at the H3K9me_3_ site. While age may be a confounding factor in mediating these changes, we explored the possibility of epigenetic reprogramming and memory due to IF. Our findings show that there is a loss of differentially expressed gene maintenance at the H3K9me_3_ locus. This may mean that both IF16 and EOD are not sufficiently robust to induce epigenetic reprogramming at this locus following refeeding, or that H3K9me_3_ may not be the sole epigenetic mechanism of IF. Therefore, the roles of other epigenetic regulatory mechanisms of IF should be consider for epigenetic maintenance.

Here we have presented a novel mechanism for how IF affects the metaboloepigenetics axis in regulating a myriad of metabolic processes to bring about robust metabolic switching responses in the cerebellum. Decoding the precise crosstalk between epigenetics and transcriptional rewiring in the context of IF, and its impact on molecular memory and other mechanisms will thus be useful in elucidating and gauging the overall benefits of IF.

## 4 Methods

### 4.1 Ethical Compliance Statement

All animal procedures were approved by the National University of Singapore Animal Care and Use Committee and performed according to the guidelines set forth by the National Advisory Committee for Laboratory Animal Research (NACLAR), Singapore. All sections of the manuscript were performed in accordance with ARRIVE (Animal Research: Reporting in Vivo Experiments) guidelines.

### 4.2 Intermittent Fasting & Refeeding Regimen

C57/BL6NTac male mice (InVivos Pte Ltd, Singapore) were raised for 3-month-old with *ad libitum* access to food using standard Teklad Global 18% protein rodent diet (Envigo, United Kingdom) and water. Mice were then randomly assigned to three study groups subjected to *ad libitum* (AL) feeding, IF for 16 hours per day (IF16), or for 24 hours on alternate days (i.e. ‘every other day’; EOD) for three months. Mice placed under IF16 regimen were provided access to food from 7 a.m. to 3 p.m. when it was removed for 16 hours daily. For EOD regimen mice, food was provided from 7 a.m. to 7 a.m. the next day, following which it was removed for the next 24 hours. All mice had *ad libitum* access to water, while the AL mice had *ad libitum* access to both food and water. In the refeeding regimen, all three groups of mice had *ad libitum* access to both food and water for two months.

During the entire experiment, the mice were housed in animal rooms at 20 to 22°C with 30 to 40% relative humidity under a 12-hour light/dark cycle. A series of physiological tests were performed on the mice. Body weight was measured weekly. Nasoanal length was measured for Lee index (body weight/nasoanal length) calculation on the day of animal euthanasia. Blood glucose and ketones were measured using a FreeStyle Optimum Meter and corresponding test strips (Abbott Laboratories, UK) at baseline and monthly via the tail bleed method. Both tests were performed at 7 a.m. Lastly, monthly food/energy consumption was recorded to measure calorie intake (weight of food consumed x kcal/g of food). The entire experimental workflow is illustrated using BioRender software **(Figure 1a)**.

### 4.3 Cerebellum Tissue Collection

Following the fasting or refeeding regimen, animals were anesthetized and euthanized. EOD mice were euthanized on a food-deprivation day. All mice were euthanized between 7 a.m. and noon. The cerebellum was harvested, immediately flash frozen and stored at −80 °C.

### 4.4 Chromatin Immunoprecipitation (ChIP)

The frozen cerebellum was crushed into a fine powder with liquid nitrogen using a pestle and mortar. The tissue was then crosslinked with 1% formaldehyde (Merck, New Jersey, United States) for 10 min at room temperature and the reaction was stopped by adding glycine (Abcam, Cambridge, United Kingdom) to a final concentration of 0.125 M for 5 min at room temperature. Fixed cells were rinsed twice with PBS with a protease (Thermo Scientific, Massachusetts, USA) inhibitor in 1:1000 ratios and resuspended in 10 ml of lysis buffer (10mM Tris-HCl pH 8, 0.25% Triton X-100, 10mM EDTA, 0.1M NaCl). The lysates were then subjected to further lysis using a Dounce tissue grinder (Sigma-Aldrich, Missouri, United States) for 14-16 strokes. The lysates were then suspended in 1% SDS lysis buffer (50mM HEPES-KOH pH 7.5, 150mM NaCl, 2mM EDTA, 1% Triton X-100, 0.1% sodium deoxycholate, 1% SDS) and ultracentrifuged at 18000rpm for 30 minutes.

Subsequently, the chromatin jelly was resuspended in 0.1% SDS lysis buffer (50mM HEPES-KOH pH 7.5, 150mM NaCl, 2mM EDTA, 1% Triton X-100, 0.1% sodium deoxycholate, 0.1% SDS) and subjected to another round of ultracentrifugation at 18000 rpm for 30 min. Next, the lysate was resuspended in 0.1% SDS buffer and sonicated for 15 cycles of 30 s on, and 30 s off in a sonicator (Diagenode, New Jersey, USA) and centrifuged at 14000 rpm for 10 min. Approximately 500 ng of sonicated chromatin was stored at −80°C as an input DNA control.

The remaining sonicated chromatin was incubated with 30 ul of Protein G Dynabeads (Invitrogen, California, United States) previously conjugated overnight to 4 ug of ChIP-grade H3K9me_3_ antibody (Abcam, Cambridge, United Kingdom). The beads were then washed for 5 minutes once, in low salt 0.1% SDS FA lysis buffer (50 mM HEPES-KOH pH 7.5, 150 mM NaCl, 2 mM EDTA, 1% Triton X-100, 0.1% sodium deoxycholate, 0.1% SDS), and then high salt 0.1% SDS FA lysis buffer (50 mM HEPES-KOH pH 7.5, 350 mM NaCl, 2 mM EDTA, 1% Triton X-100, 0.1% sodium deoxycholate, 0.1% SDS), followed by a LiCl buffer wash (10 mM Tris-HCl pH 8, 0.25 mM LiCl, 1 mM EDTA, 0.5% NP40, 0.5% sodium deoxycholate). Lastly, the immunoprecipitated material was washed twice with TE buffer (10 mM Tris-HCl pH 8, 1 mM EDTA), and elutedwith ChIP elution buffer (50 mM Tris-HCl pH 8, 10 mM EDTA, 1% SDS) incubation for 2 hours at 68°C for both input and immunoprecipitated DNA. Finally, chromatin was incubated for 1-hour with RNase at 37°C, and digested with 10 ul of 5 mM NaCl and 5 ul of Proteinase K overnight at 68°C. DNA was then extracted using phenol-chloroform and eluted in nuclease-free water.

Extracted DNA was quantified using Quantifluor double-stranded DNA kit (Promega Corporation, Australia) as per manufacturer instruction. 20X TE buffer was first diluted 20 times, before diluting the Quantifluor dsDNA dye in 1:400 ratio to prepare the working solution. In each sample, 1 ul of DNA was diluted in 200 ul of working solution, whereas for blank and standard, 2 ul of 1xTE buffer and 2 ul of provided DNA standard (100 ng/ul) were diluted in similar volume of working solution. Both blank and standard solution were used for calibration to plot a standard curve before actual measurement of DNA concentration. For fragment size analysis, 100 ng of DNA was first purified using a QIAquick PCR purification kit (Qiagen, Japan) according to the manufacturers’ protocol. The resulting purified DNA was then mixed with a loading dye (Thermo Scientific, Massachusetts, USA) at a 1:50 ratio and run in a 1% agarose gel (Lonza, Switzerland) at 100 V for 30 min and stained with SYBR safe dye (Invitrogen, California, United States). Imaging was carried out using ChemiDocXRS+ imaging system (Bio-Rad Laboratories, California, USA). Fragments of the ideal size of 100-500 base-pairs were selected for the analysis.

A library was prepared using New England Biolabs Ultra II DNA Kit for Illumina (New England Biolabs, United States) according to the manufacturer’s instructions. The immunoprecipitated material was sequenced using the 150-bp paired-end protocol provided by Illumina Genome Analyzer 1.9 (Novogene, Beijing). All data obtained from each sample were pooled for analysis. A brief experimental workflow is illustrated using BioRender software **(Figure 1b)**.

### 4.5 ChIP Sequencing Bioinformatics Analysis

Raw reads were checked for quality using FastQC (Babraham Bioinformatics, United Kingdom). Reads with low quality (proportion of low-quality bases larger than 50%) or N ratio (unsure base) larger than 15% were discarded. Moreover, reads with adaptor at the 5’-end or those without adaptor and inserted fragment at the 3’-end were also discarded as well. Next, the reads will be trimmed at the adaptor sequence at the 3’-end. A further discarding of reads whose length are less than 18 units of nucleotides following trimming will be conducted. The reference genome for Mus musculus (mm10) and gene model annotation files were downloaded from the National Center for Biotechnology Information (NCBI) genome database. Reference genome indexing and mapping of the quality checked paired-end reads to the reference genome was carried out using the Burrows-Wheeler Alignment Tool (v0.7.17) (Li and Durbin, 2010). MACS2 callpeak and MACS2 bdgdiff software (Zhang et al., 2008) peak detection and comparisons, respectively. Strand cross-correlation was computed on the resulting peaks and plotted using GraphPad Prism software (v5). Heatmaps, fingerprint, and principal component analysis (PCA) plots were generated using DeepTools2 software (Ramírez et al., 2016). Three-dimensional PCA plot was constructed using Sigmaplot software (v1.3). Venn diagrams were prepared using Bioinformatics & Evolutionary Genomics online software (Bioinformatics & Evolutionary Genomics, Belgium). The functional significance of peaks was analyzed using webserver g:Profiler (Reimand et al., 2016) and DAVID (Jiao et al., 2012). Gene ontology (GO) terms with adjusted p-value <0.05 were considered as significantly enriched among the pool of differentially expressed peaks. Diagrams representing GO enrichment analysis results were plotted using GraphPad Prism software (v5) and ggplot2 R package (v3.1.1).

### 4.6 Total Eukaryotic mRNA Extraction

RNA from cerebellum tissue was isolated using EZ-10 DNAaway RNA Mini-Preps Kit (Bio Basic, Canada) according to the manufacturers’ protocol. Briefly, frozen cerebellum samples were homogenized and lysed in the lysis buffer. Contamination of genomic DNA was prevented by using the gDNA eliminator column. Purity of RNA was determined by using the Nanodrop ND-1000 (Thermo Fisher Scientific, USA), while RNA integrity was assessed by agarose gel electrophoresis as well as the Agilent 2100 Bioanalyzer (Agilent, USA). Enriched RNA was of high-quality, demonstrating an OD_260_/OD_280_ ratio of 1.9-2.0 from Nanodrop readings, two distinct bands indicating 28S and 18S following agarose gel electrophoresis, and RNA integrity number ≥ 6.8 with a smooth base line using the Agilent 2100 Bioanalyzer.

Following the isolation of high-quality and pure total RNA from cerebellum tissues, cDNA library was constructed using the NEBNext^®^ Ultra^TM^ RNA library preparation kit as per the manufacturers’ protocol (New England BioLabs, USA). mRNA was first purified via the addition of poly-T-oligo-attached magnetic beads, and subjected to random fragmentation using a fragmentation buffer. The first strand of cDNA was synthesized using a random hexamer primer and RNase H-(M-MuLV reverse transcriptase). The second strand of cDNA was synthesized using DNA polymerase I and RNase H, and the resulting double stranded cDNA was then purified using AMPure XP beads. The overhangs of these purified double-stranded cDNA were further processed by exonuclease and polymerase to create blunt ends, and the 3’ ends of these DNA fragments were adenylated and subsequently ligated with the NEBNext hairpin loop structure adaptor on both ends for hybridization. For optimal isolation of cDNA fragments of approximately 150-200 base pairs in length, the DNA fragments were purified using the AMPure XP system (Beckman Coulter, USA), followed by PCR amplification and purification by AMPure XP beads to obtain the DNA fragments representing the complete library. The resulting libraries were sequenced using HiSeq^TM^ 2500 Illumina platform to obtain a minimum 12GB raw data per sample (Illumina, USA). A brief experimental workflow was illustrated using BioRender software **(Figure 1b)**.

### 4.7 mRNA Sequencing Bioinformatics Analysis

The reference genome for *Mus musculus* (mm10) and gene model annotation files were downloaded from the National Center for Biotechnology Information (NCBI) genome database. The reference genome was indexed and the paired-end quality checked reads were mapped to the reference genome using the STAR aligner (v2.5) (Dobin et al., 2013). Reads mappingto each gene were qualified using HTSeq (v0.6.1) (Anders et al., 2015). Fragments per kilobase of exon model per million mapped reads (FPKM) for each gene were computed based on the gene length and the number of reads mapped to the gene. The FPKM value was then used for estimation of gene expression levels. A total of 35275 unique RNA transcripts were quantified in the cerebellum datasets. Differential gene expression analysis was performed using the DESeq2 R Package (v2_1.6.3) (Anders and Huber, 2010) Using a negative binomial distribution model for the gene counts. The resultant p-values were then adjusted using Benjamini and Hochberg’s test to control false discovery rate (FDR). Genes with adjusted p<0.05 were assigned as differentially expressed.

PLS-DA (Partial least squares-discriminant analysis) plots were constructed using the mixOmics R package (v6.6.2) (Rohart et al., 2017). Venn diagrams were prepared using online software (Bioinformatics & Evolutionary Genomics, Belgium). Heatmaps were built using pheatmap R package (v1.0.12) (Rohart et al., 2017), while the volcano plots in this study were prepared using the ggplot2 R package (v3.0.0). Gene ontology (GO) enrichment analysis for differentially expressed genes was carried out using g:Profiler (Reimand et al., 2016), DAVID (Jiao et al., 2012) and Enrichr (Kuleshov et al., 2016) webservers. GO terms with adjusted p-value < 0.05 were considered to be significantly enriched among the pool of differentially expressed genes. Enrichment analysis diagrams were plotted using GraphPad Prism software (v5) and ggplot2 R package (v3.1.1).

### 4.8 Integrative ChIP and mRNA Sequencing Analysis

To identify the number of differential expressed genes that were modulated by H3K9me_3_, a Venn analysis was performed (Bioinformatics & Evolutionary Genomics, Belgium) and illustrated using Inkscape software (v0.92) as Euler diagrams. Next, we performed a series of GO interrogation and visualization of the overlapped genes using GOplot R package (v1.0.2) (Walter et al., 2015) and Inkscape software (v0.92) respectively. ChIP peaks and mRNA tracks were visualized in the Integrative Genome Browser (v2.5.3) (Robinson, 2012; Robinson et al., 2017; Thorvaldsdóttir et al., 2013). Network analysis and visualization of the overlapped genes were achieved using the STRING (v11.0) (Szklarczyk et al., 2019) webserver and Cytoscape software (v3.7.1) (Paul Shannon et al., 2003). The ClusterOne (v1.0) plugin in Cytoscape was used with default parameters to obtain significant gene clusters. Metabolic pathways analysis was painted as metabolic maps using iPath3 online software tool (v3) (Darzi et al., 2018) and metabolic cellular functions performed using BioCyc Omics dashboard software (19.0) (Karp et al., 2016; Paley et al., 2017).

### 4.9 Statistical Analysis

Experimental data were analyzed using GraphPad Prism software (v5) for statistical analysis. Two-way analysis of variance (ANOVA) was used, followed by Dunnett’s post-hoc test, for weight, glucose and ketone measurement. Numerical values were expressed as mean ± standard error of the mean (S.E.M). A p-value<0.05 was considered statistically significant. Correlation was determined using the “cor.test” function in R with options set alternative= “greater” and method= “Spearman”. All experiments were performed using at least three biological replicates per condition, and Pearson correlation coefficients of at least 0.9 demonstrated high coverage and reproducibility **(Figure S3)**. Statistical analysis for bioinformatics was performed associated documentation default parameters.

## Accession Numbers

High-throughput sequencing data have been submitted to the NCBI Sequence Read Archive (SRA) under accession number GSE135945.

## Supplemental Information

Supplemental Information includes 08 figures, 05 tables and can be found with this article online.

## Acknowledgements

We thank Novogene (Beijing, China) for their kind assistance in data processing. The Singapore National Medical Research Council Research Grants (NMRC-CBRG-0102/2016 and NMRC-OFIRG-036/2017) supported this work. This study also supported by the National Research Foundation (NRF) funded by the Korean Government (NRF-2019R1A2C3011422 and NRF-2019R1A5A2027340)

