## Supplementary Figure Legends for "Integrative Epigenomic and Transcriptomic Analysis Reveals Robust Metabolic Switching in the Brain During Intermittent Fasting"

**SUPPLEMENTAL INFORMATION**

**Supplemental Figures and Figure Legends.**

**Figure S1: IF protects mice from weight gain without corresponding changes in caloric intake.**

**(a)** IF mice have significantly lower body weight than AL mice after three-months of IF. No significant body weight change is observed between IF16 and EOD mice. Following two-months refeeding, no significant average body weight change is observed between biological groups. **(b-c)** No significant difference in the overall energy intake or compositional intake is observed across mice subjected to IF or refeeding as compared to control. Data are represented as mean ± S.E.M; One or Two-way ANOVA with Tukey’s and Dunnett’s *post-hoc* analysis respectively. ****p<0.0001 vs AL.

**Figure S2: IF initiated metabolic switching in a temporal-dependent manner.**

**(a-b)** Significant lowering of blood glucose and higher blood ketone levels shown in both IF16 and EOD mice, as compared to AL. Following refeeding, mice do not display significant differences in their blood glucose and ketone level across biological groups. The insets demonstrate total average blood glucose and ketone level over time. **(c)** Body composition analysis demonstrates significant reduction in Lee index and body fat mass in both IF16 and EOD mice, as compared to AL. Significant difference correlates with increasing time-restricted feeding period. Data are represented as mean ± S.E.M; One or Two-way ANOVA with Dunnett’s *post-hoc* analysis. *p < 0.05, **p < 0.005, ***p < 0.001 and ****p<0.0001 vs AL. AUC, area under curve.

**Figure S3: Pearson correlation coefficients for IF and refeeding epigenomic and transcriptomic dataset.**

Pearson correlation coefficients of minimal 0.9 value obtained for epigenomic and transcriptome samples demonstrated high coverage and reproducibility.

Figure S4: Comparative analysis of H3K9me_3_-dependent regulation of differentially expressed peaks functional annotation network as a result of IF and refeeding.

(a-b) Differentially expressed peaks as a result of IF and refeeding are used as input and clusters were identified with ClusterOne highlighted in different colors. Functional annotation of each clusters is labelled beside. Differentially expressed peaks that are maintained between IF and refeeding are labelled as black and their clusters and functional ontologies are shown in the middle of the diagram surrounded by a rectangle. Each dot represents a gene symbol.

Figure S5: Comparative analysis of differentially expressed genes functional annotation network as a result of IF and refeeding.

(a-b) Differentially expressed genes as a result of IF and refeeding are used as input and clusters were identified with ClusterOne highlighted in different colors. Functional annotation of each clusters is labelled beside. Differentially expressed genes that are maintained between IF and refeeding are labelled as black and their clusters and functional ontologies are shown in the middle of the diagram surrounded by a rectangle. Each dot represents a gene symbol.

Figure S6: Bar chart illustrating pathways enrichment as a result of IF.

(a-b) H3K9me_3_-controlled differentially expressed genes during IF16 and EOD are input into Enrichr to obtain pathways enrichment. Significant H3K9me_3_-controlled differentially expressed gene involved in pathway enrichment of IF16 and EOD compared to control plotted against statistical significance (represented as (-log_10_ p-value). Downregulated pathways are shown on the left whereas upregulated pathways are shown on the right.

Figure S7: Global metabolic map for H3K9me_3_-controlled differentially expressed genes during IF.

(a-b) Metabolic map showing the myriad of pathways and enzyme-related modules exhibited by H3K9me_3_-controlled differentially expressed genes during IF16 and EOD. The complexity and pattern of metabolic pathways differ between both IF16 and EOD. Pathways and enzyme modules are highlighted in red.

**Figure S8: Metabolic subsystems of** H3K9me_3_-dependent regulation of differentially expressed genes.

(a-b) H3K9me_3_-dependent regulation of differentially expressed genes during IF16 and EOD are input into BioCyc omics dashboard. A system of cellular metabolic functions, namely biosynthesis and degradation, is plotted on the x-axis whereas relative expression level is plotted as log (fold-change) in the y-axis.

**Supplemental Table Legends.**

Table S1: Enrichment Analysis of H3K9me_3_ Differential Peaks for Intermittent Fasting & Refeeding.

Table S2: Enrichment Analysis of Differentially Expressed Genes for Intermittent Fasting & Refeeding.

Table S3: Enrichment Analysis of H3K9me_3_ Epigenomic & Transcriptomic Age Associated Changes.

Table S4: Enrichment Analysis of H3K9me_3_-Dependent Transcriptomic Changes in Metabolism during IF16 & EOD.

Table S5: Pathway Enrichment Analysis of H3K9me_3_-Dependent Transcriptomic Changes in Metabolism during IF16 & EOD.
