## Supplementary figures and images for "Integrative Epigenomic and Transcriptomic Analysis Reveals Robust Metabolic Switching in the Brain During Intermittent Fasting"

### Supplementary Figure 1

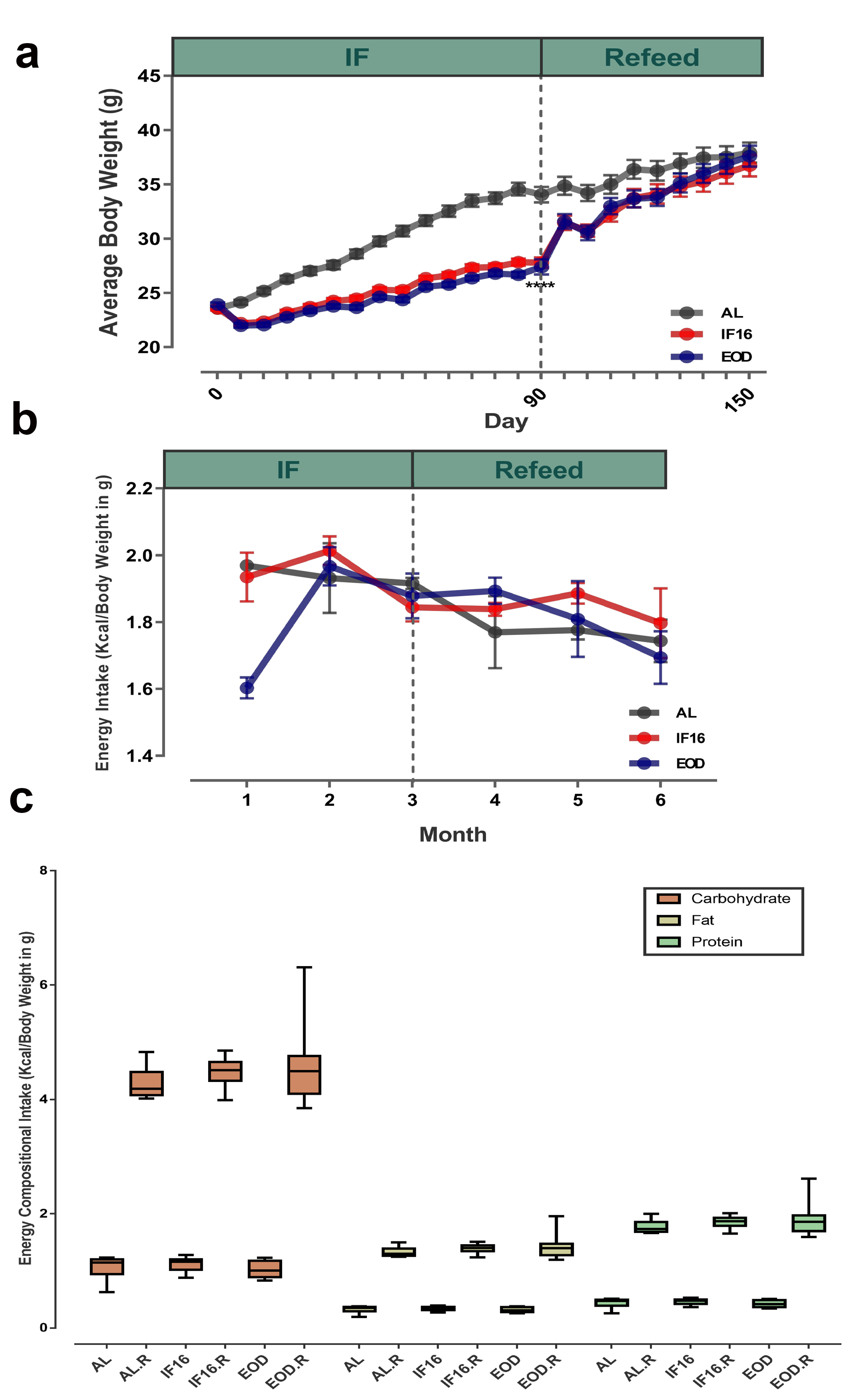

### Supplementary Figure 2

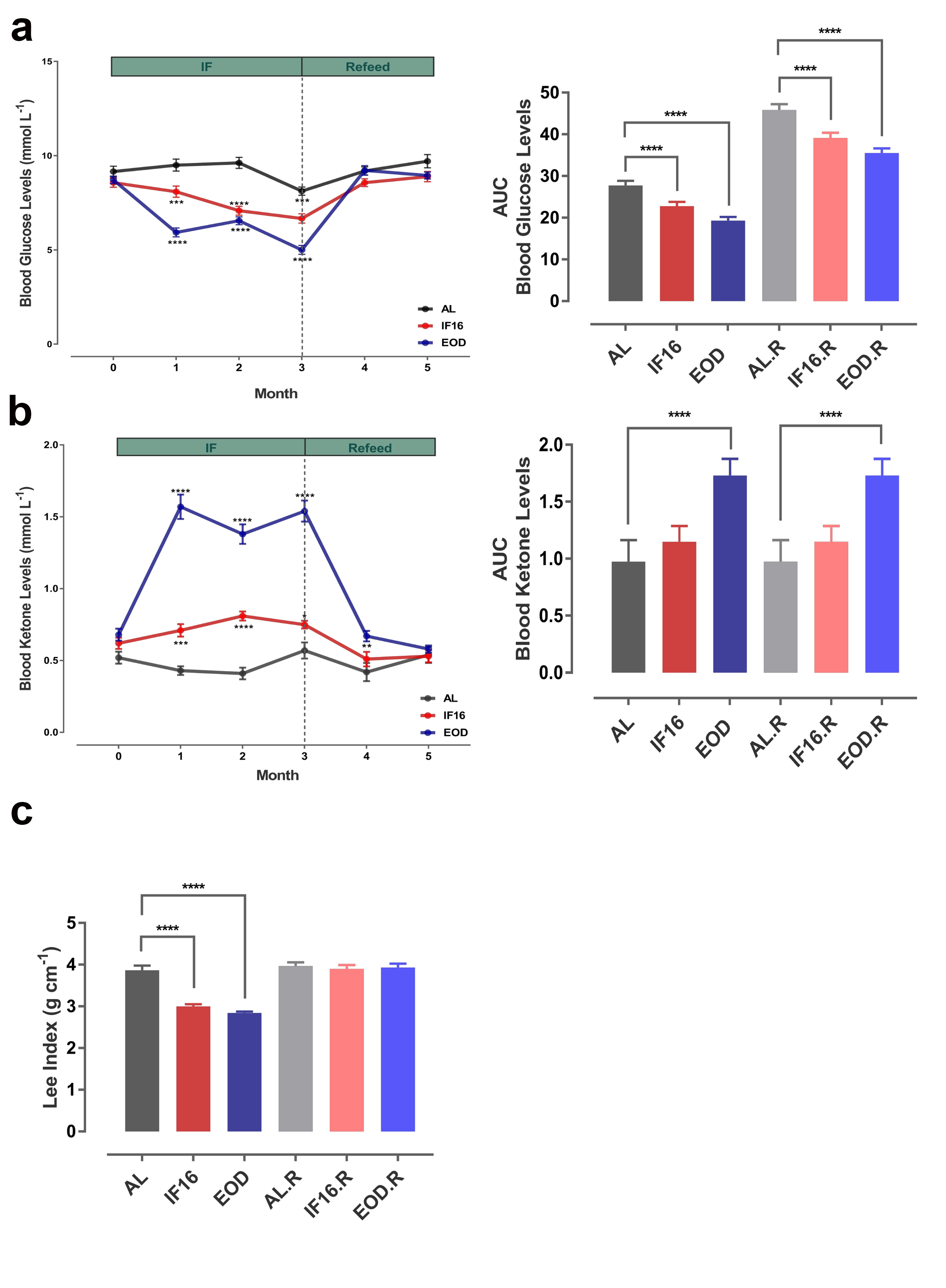

### Supplementary Figure 3

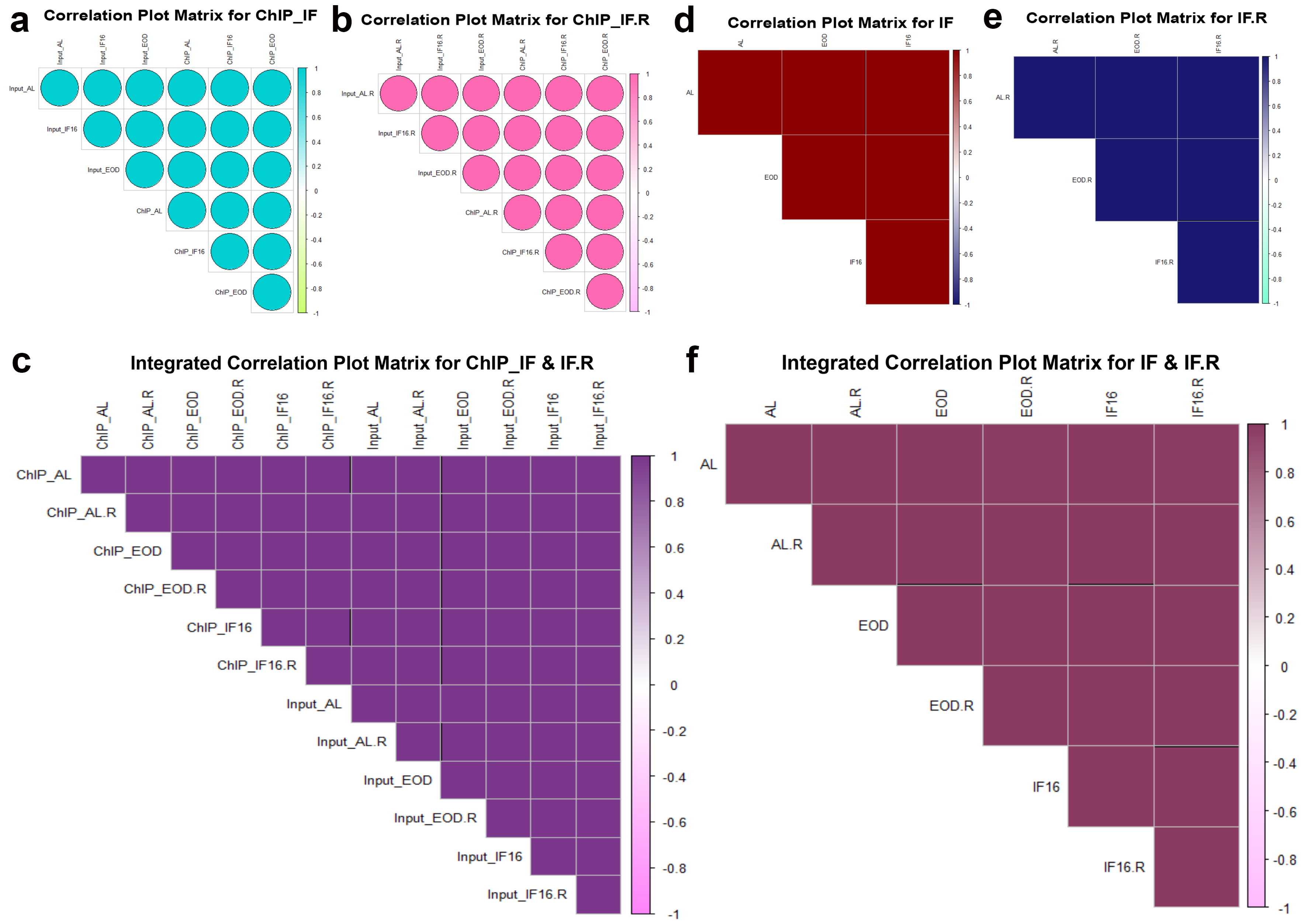

### Supplementary Figure 4

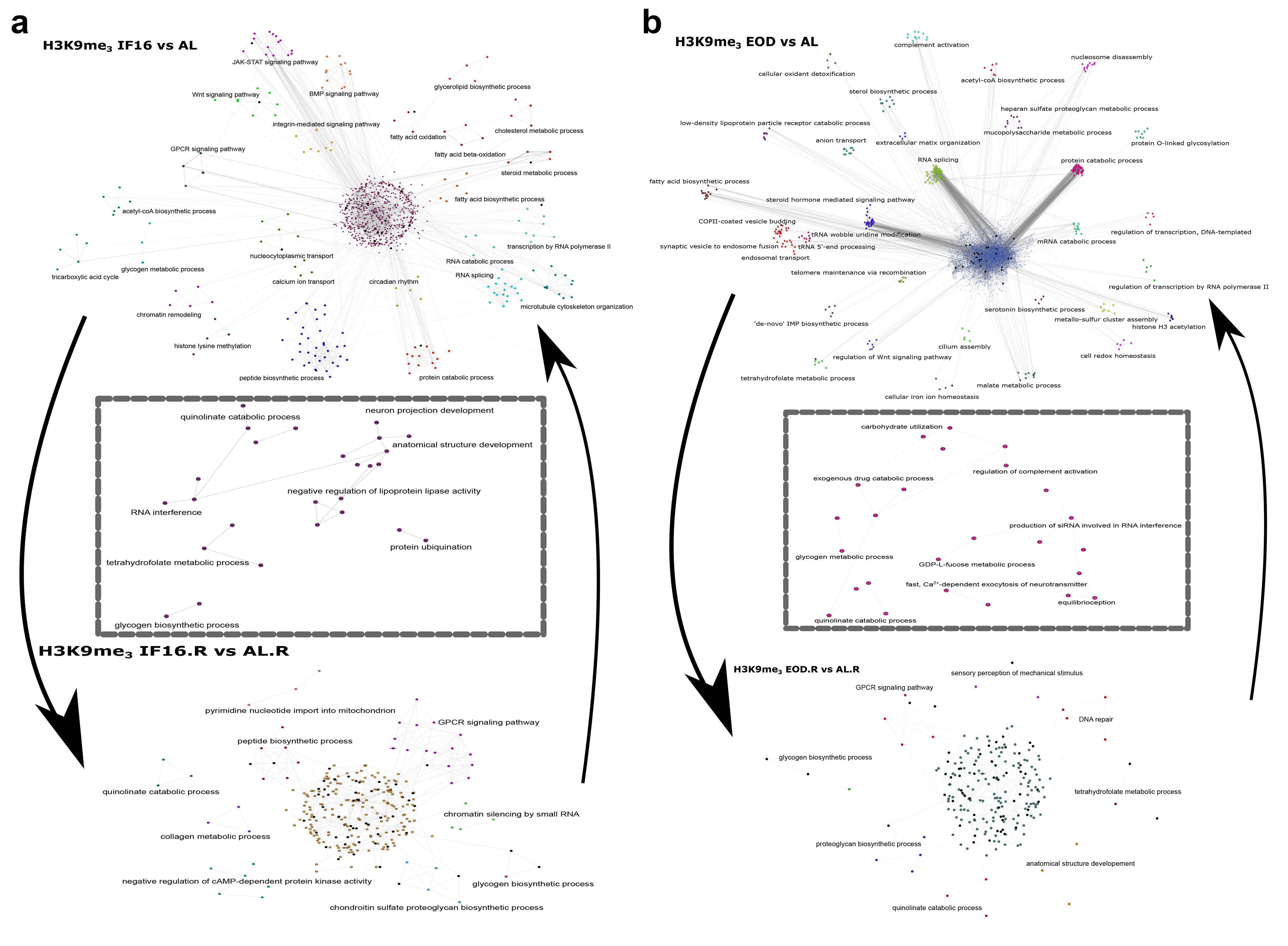

### Supplementary Figure 5

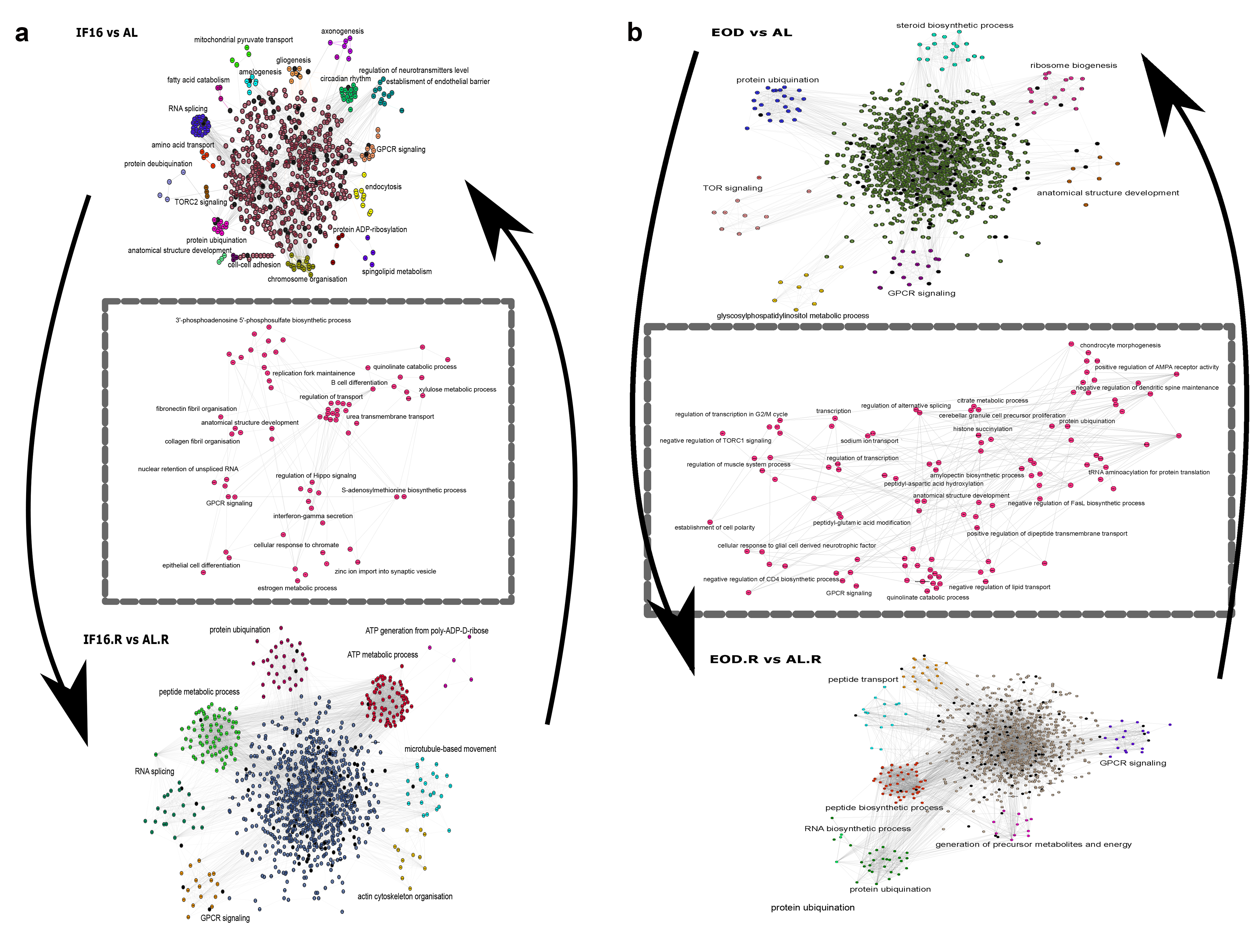

### Supplementary Figure 6

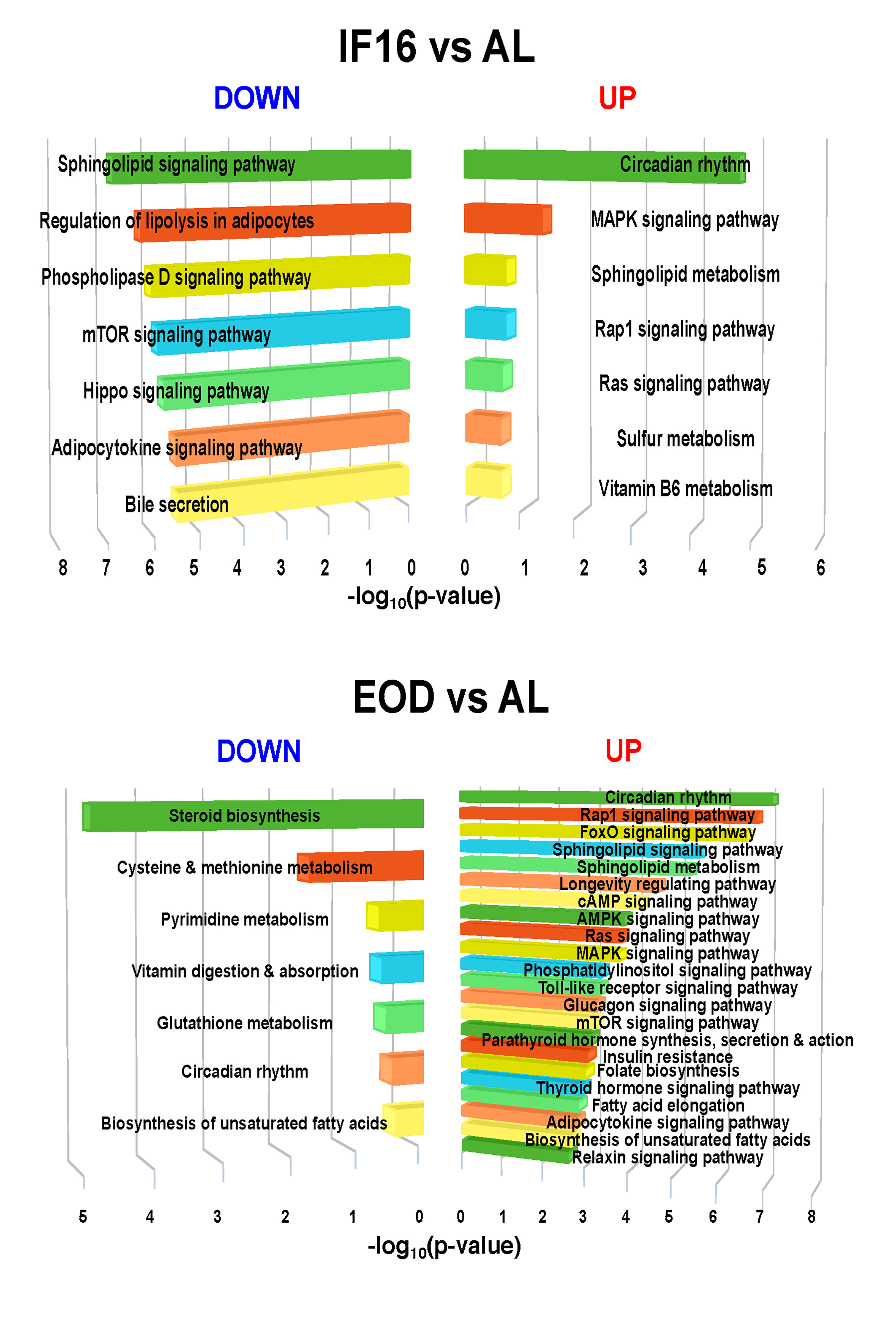

### Supplementary Figure 7

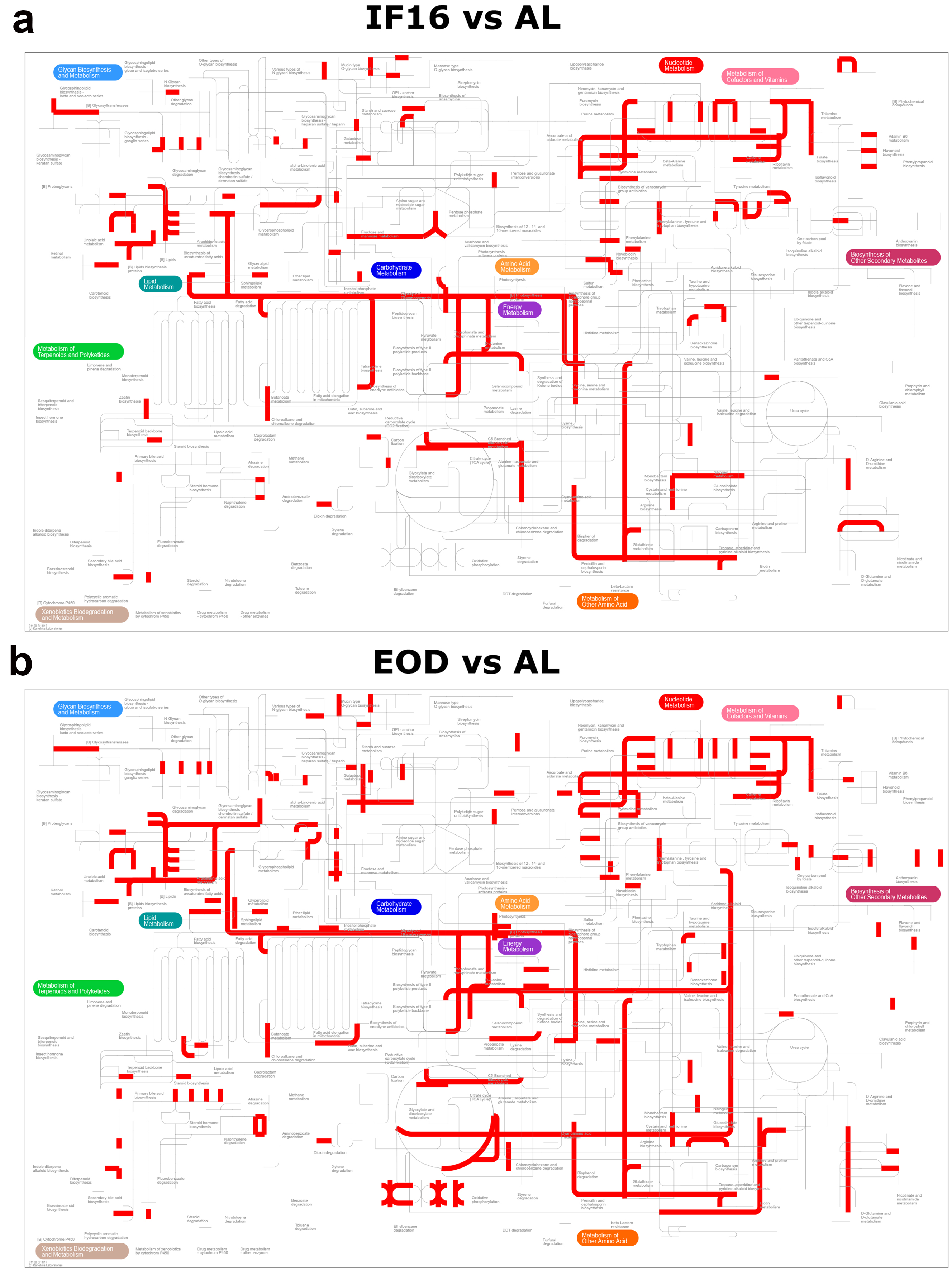

### Supplementary Figure 8

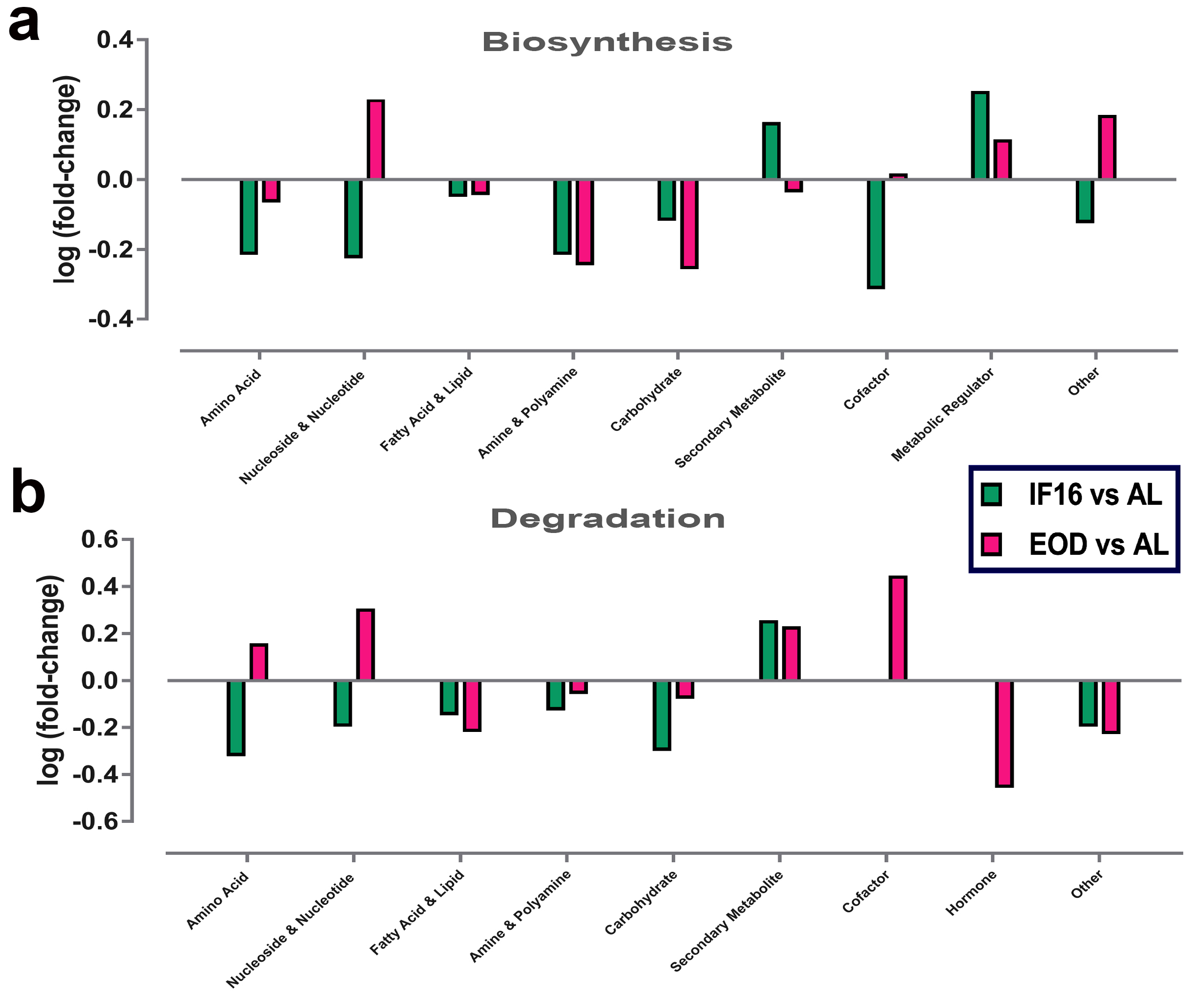
